# Zebrin molecular identity and cerebellar location determine Purkinje cell vulnerability in Christianson syndrome mice

**DOI:** 10.64898/2026.01.05.697573

**Authors:** Louis-Charles Masson, Julia Tourbina-Kolomiets, Brenda Toscano Marquez, Alanna J. Watt, R. Anne McKinney

**Author notes:** Corresponding senior authors. Joint Contribution (Louis-Charles Masson, Julia Tourbina-Kolomiets).

## Abstract

Ataxia is a major characteristic feature in Christianson syndrome, where Purkinje cells in the anterior cerebellar vermis are vulnerable to degeneration while those in the posterior vermis are resilient. Here, we provide a temporal study of changes in Purkinje cell function, cell vulnerability and innervation of cerebellar nuclei in a mouse model of Christianson syndrome. Purkinje cells express certain molecules in a patterned manner across the cerebellum, such as aldolase C/zebrin-II (zebrin). We show that zebrin patterning appears normal before disease onset in Christianson syndrome mice with no apparent cell death, similar to what has been reported in patients. We observe rapid Purkinje cell death in the anterior lobe of Christianson syndrome mice that is exclusive to zebrin-negative Purkinje cells, with zebrin-positive Purkinje cells showing resilience. Intrinsic firing deficits exclusively in the anterior cerebellum are observed at the same time as the first observed cell death in the anterior cerebellum. In contrast, posterior Purkinje cells degenerated later, but zebrin-negative and zebrin-positive cells were equally susceptible to degeneration. Purkinje cells’ innervation in the cerebellar nuclei was also lost in Christianson syndrome mice in a region-specific manner: neurons in the anterior cerebellar nuclei, with predominantly zebrin-negative Purkinje cells’ inputs, displayed dramatic loss of Purkinje cells’ innervation, while neurons in the posterior cerebellar nuclei, innervated by both zebrin-negative and zebrin-positive terminals, showed greater resiliency. Together, our results highlight that cerebellar location and zebrin molecular identity appear important to the vulnerability of Purkinje cells and their innervation in the cerebellar nuclei in Christianson syndrome.

**HIGHLIGHTS:**

- First study showing Purkinje cell intrinsic firing alterations in Christianson syndrome mice.
- Zebrin patterning appears normal before disease onset in Christianson syndrome mice.
- Purkinje cell firing deficits and death start at post-natal day 35 only in the anterior cerebellum in a Christianson syndrome mouse model.
- Zebrin-negative and zebrin-positive Purkinje cells in the posterior cerebellum are initially resilient but are both equally susceptible to Purkinje cell death at later stages of the disorder.
- Zebrin-negative Purkinje cell inputs to the anterior cerebellar nuclei are lost, while zebrin-positive inputs remain.
- The pattern of Purkinje cell death in Christianson Syndrome is similar to that of many rare ataxias, suggesting a comparable underlying pathophysiology, which could lead to common therapeutic targets.

## 1. INTRODUCTION

Christianson syndrome (CS) is a rare X-linked neurodevelopmental and neurodegenerative disorder characterized predominantly by non-verbalism, ataxia, microcephaly, cerebellar atrophy, epilepsy, and severe intellectual disability (Christianson et al., 1999; Kavanaugh et al., 2024; Schroer et al., 2010; Strømme et al., 2011). Males are predominantly affected, with females exhibiting milder, usually more psychiatric symptoms (Christianson et al., 1999; Pescosolido et al., 2014; Schroer et al., 2010; Strømme et al., 2011). Gilfillan *et al*. (2008) (Gilfillan et al., 2008) discovered a common feature shared among all patients, first reported in a South African pedigree by Christianson *et al*. (Christianson et al., 1999): mutations in the *SLC9A6* gene encoding the endosomal sodium-hydrogen exchanger 6 (NHE6) (Gilfillan et al., 2008; Pescosolido et al., 2014; Strømme et al., 2011; Toscano Márquez et al., 2021). To date, over 100 different mutations in the *SLC9A6* gene have been discovered, resulting in the loss of functional NHE6 protein (Gilfillan et al., 2008; Schroer et al., 2010). Most CS patients exhibit severe truncal and/or gait ataxia, a debilitating condition characterized by motor coordination and balance deficits, caused by cerebellar dysfunction (Amore et al., 2021; Kavanaugh et al., 2024; Xu et al., 2017).

Patterned death of Purkinje cells (PCs), essential inhibitory neurons acting as the sole output of the cerebellar cortex, is a common feature observed in many ataxias (Gomez et al., 1997; Sarna & Hawkes, 2003; Toscano Márquez et al., 2021; Xu et al., 2017). In CS, PC degeneration was previously reported to be selectively in PCs in the anterior region (Strømme et al., 2011; Xu et al., 2017), a pattern which is strikingly similar to that observed in many other rare ataxias, including Autosomal-Recessive Spastic Ataxia of Charlevoix-Saguenay (ARSACS) (Toscano Márquez et al., 2021) and spinocerebellar ataxia type 6 (SCA6) (Gierga et al., 2009; Yang et al., 2000) and in aging (Donofrio et al., 2025). To better understand patterned PC vulnerability in ataxia, recent research efforts have focused on PC molecular identity (Donofrio et al., 2025; Toscano Márquez et al., 2021). PCs are known to express specific molecules in a patterned way across cerebellar regions (Voogd, Jaarsma, & Marani, 1996), perhaps the most notable being aldolase C, also known as zebrin-II (zebrin) (Hawkes & Herrup, 1995; Toscano Márquez et al., 2021). Zebrin is a brain-specific glycolytic enzyme (Hawkes & Herrup, 1995), expressed in parasagittal stripes across the cerebellum in a pattern that matures by postnatal day (P) 24 in rodents, is highly reproducible between individuals and species, and is maintained for life (Hawkes & Herrup, 1995; Lin et al., 2020). Many models of ataxia exhibit zebrin-negative PC vulnerability (Hawkes & Herrup, 1995; Sarna & Hawkes, 2003), raising the possibility of zebrin-positive PC identity being neuroprotective (Slemmer et al., 2007).

Studies on CS ataxia have been limited; anterior PC vulnerability has previously been reported in the CS murine model at age P57-P60, affecting zebrin-negative PCs preferentially (Strømme et al., 2011; Xu et al., 2017). Moreover, it has also been shown that PCs in the central and nodular lobes, which are mostly zebrin-positive, are most resistant to degeneration in CS (Strømme et al., 2011; Xu et al., 2017). However, important information remains unknown regarding the development of zebrin bands and the timeline of zebrin-negative and zebrin-positive PC degeneration in CS, especially in the posterior cerebellum. Thus, in this study, we sought to establish a timeline of zebrin-related selective neuronal vulnerability in the CS murine cerebella, comparing PCs of different zebrin molecular identities in motor-associated anterior lobule III and posterior lobule VIII/IX (Lin et al., 2020; Stoodley, Valera, & Schmahmann, 2012), to determine more precisely which subset of PCs degenerate and when.

As the cerebellar nuclei (CN) are the final integration center of cerebellar information, receiving strong inhibitory input from PC axons, reduced innervation onto the CN can strongly impact cerebellar function and motor control. Although more research is needed to elucidate how PCs from different lobules integrate onto the CN, we know that zebrin-negative and zebrin-positive PCs target different areas of the CN (Sugihara, 2011). Interestingly, in ARSACS, which shares a similar pattern of PC vulnerability with CS (Larivière et al., 2015; Strømme et al., 2011), a decreased connectivity to the anterior CN region was observed with zebrin-negative PCs decreasing their inputs onto the CN, while zebrin-positive cells did not (Toscano Márquez et al., 2021). While previous work has shown that anterior PCs are more vulnerable to degeneration in CS (Sikora et al., 2016; Strømme et al., 2011), how PC loss affects connectivity in the CN in CS remains unknown. Here, we sought to investigate how cerebellar cortex PC loss affects CN innervation in CS, comparing zebrin-related innervation in anterior and posterior regions of the CN over time.

Although the exact role of zebrin identity and parasagittal compartmentalization remains to be fully elucidated, recent research efforts have highlighted potential mechanisms involved. In the absence of synaptic input, PCs continue to fire spontaneous action potentials with high frequency and regularity and are thus often referred to as pacemaker cells (Häusser & Clark, 1997; Masoli, Solinas, & D’Angelo, 2015; Raman & Bean, 1997; Watt et al., 2009). The spiking activity of PCs is of critical interest, as PC firing deficits often present as the first sign of cerebellar dysfunction, preceding PC loss (Ady et al., 2018; Cook, Fields, & Watt, 2021). We know that PC firing properties are greatly influenced by zebrin identity (Zhou et al., 2014). Notably, zebrin-negative PCs, which are more prevalent in anterior lobules (Sugihara & Shinoda, 2004), fire action potentials with higher frequency than their zebrin-positive counterparts, which are more prominent in posterior lobules (Sugihara & Shinoda, 2004; Zhou et al., 2014). Some have hypothesized that the higher spiking rate of zebrin-negative PCs could play a role in their selective vulnerability observed in several models of ataxia, due to the greater energy demand (De Zeeuw & Ten Brinke, 2015; Howarth, Gleeson, & Attwell, 2012). As PC intrinsic firing has been shown to be a promising therapeutic target in ataxia (Cook, Fields, & Watt, 2021), we sought to investigate PC firing patterns in CS over time.

We found that PC intrinsic firing deficits occur only in anterior PCs at a similar time as the onset of cell death, at P35. Further, only anterior zebrin-negative PCs are degenerating at this early-disease stage, while anterior zebrin-positive PCs and all PCs in the posterior are resilient. Similarly, only zebrin-negative PC innervation onto the anterior CN is reduced in CS at this stage. At later disease stages, starting at P150, zebrin-positive PCs in the anterior and PCs in the posterior, independently of zebrin identity, degenerate as well. Surprisingly, PC input to the posterior CN was only found to be reduced at P150 but not P350 and the reduction was specifically in zebrin-positive PC innervation. Altogether, these findings support the hypothesis that both molecular location and cerebellar identity influence PC vulnerability in CS.

## 2. METHODS

### Transgenic animals

The *Slc9a6* knockout mice, Slc9a6^tm1Dgen^ mice, bred on a C57/BL6 background and deficient in the *Nhe6* gene were purchased from Jackson Laboratories and used as previously described (Strømme et al., 2011). Briefly, these mice were engineered by inserting a LacZ-Neo cassette into exon 6 of the *Slc9a6* gene, introducing a stop-codon and a polyadenylation termination signal, causing significantly reduced transcription of the Nhe6 transcript following the LacZ-Neo cassette. KO males (herein referred to as CS mice) were used in the experiments described in the present study, with age-matched C57/BL6 males (WT mice) purchased from Jackson Laboratories used as controls. Only male mice were used in this study due to CS being an X-linked disorder (Christianson et al., 1999) and females having milder symptoms, with many not exhibiting ataxia. Experiments were performed at ages P25, P35, P50, P150 and P350. A minimum of three mice of each genotype were used in each experiment. All animal handling procedures were carried out according to the guidelines of the Canadian Council on Animal Care and the National Research Council’s Guide for the Care and Use of Laboratory Animals and approved by the McGill University Animal Care Committee. The animals were maintained by Tanya Koch at the McGill University Comparative Medicine and Animal Resources.

### Ex vivo acute slice electrophysiology

Acute cerebellar slices were prepared *ex vivo* as previously described (Ady et al., 2018). Mice were deeply anesthetized with isoflurane and rapidly decapitated. Brains were removed and placed in ice-cold 95% O_2_ and 5% CO_2_ saturated artificial cerebrospinal fluid (ACSF). ACSF contained: 125 mM NaCl, 2.5 mM KCl, 1 mM MgCl_2_, 1.25 mM NaH_2_PO_4_, 25 mM NaHCO_3_, 2 mM CaCl_2_, and 25 mM glucose, with a final osmolality of 320 mOsm and pH 7.4. *Ex vivo* acute sagittal slices of 200 µm thickness were cut from the cerebellar vermis using a VT1200S microtome (Leica Microsystems, Wetzlar, Germany). Slices were incubated for 30-45 min at 37°C and then placed at room temperature for up to an additional 8 h in continuously bubbling ACSF prior to further analysis. Spontaneous intrinsic PC firing was non-invasively recorded at 33.5-34.5°C using extracellular loose patch recording with ACSF-filled electrodes pulled using a P-1000 puller (Sutter Instruments, Novato, CA, USA), as previously described (Ady et al., 2018). Recordings were done with the application of a cocktail of inhibitors of fast excitatory and inhibitory synaptic inputs to Purkinje cells, using 100μM picrotoxin, 10 μM NBQX and 50μM AP-V. Firing frequency and coefficient of variation (CV) were analyzed with Clampfit 11.4 (San Jose, California, United States). Anterior lobule recordings were made in lobule III, while posterior lobule recordings were made in lobule VIII.

### Tissue preparation for immunohistochemistry

WT and CS mice were deeply anesthetized with isoflurane prior to being sacrificed. At P25, brains were extracted and placed in 4% paraformaldehyde (PFA) for 5 days at 4°C, followed by washes with 0.1 M phosphate buffer (PB, pH 7.4), and transferred to 0.1 M PB containing 0.05% sodium azide for storage at 4°C. For all other timepoints, mice were perfused transcardially, under deep anesthesia with isoflurane, with ice-cold phosphate-buffered saline (PBS, pH 7.4), followed by transcardial fixation perfusion using ice-cold 4% PFA. Brains were then extracted, and immersion fixed overnight in 4% PFA at 4°C, before being washed with 0.1 M PB, and transferred to 0.1 M PB containing 0.05% sodium azide for storage at 4°C.

### Immunohistochemistry

Immunohistochemistry was performed on coronal and sagittal free-floating cerebellar slices of 100 µm thickness obtained using a 5100mz vibratome (Campden Instruments, Lafayette, IN, USA). Slices were incubated overnight in permeabilization solution (0.1 M PB, 1.5% heat-inactivated horse serum, 0.4% triton X-100, pH 7.4), then incubated with primary antibody diluted in permeabilization solution for 3 days at 4°C on a 60rpm shaker. Slices were then washed with blocking buffer (0.1 M PB, 1.5% heat-inactivated horse serum, pH 7.4), then incubated with secondary antibody diluted in blocking buffer for 2 hrs at room temperature on a 60rpm shaker, then washed with 0.1 M PB and mounted on Superfrost plus microscope slides (Fisher Scientific, Pittsburg, PA, USA) using Dako fluorescent mounting medium (Agilent Technologies, Santa Clara, CA, USA).

### Cerebellar cortex

Cerebellar coronal slices were stained with primary antibodies: goat anti-Aldolase C (1:500, N-14; Santa-Cruz Biotechnology, Dallas, TX, USA) and rabbit anti-calbindin (1:1000, CB38A; Swant, Marly, Switzerland). Primary antibodies were then tagged to secondary antibodies: Alexa-488 conjugated donkey anti-goat (1:250; Invitrogen, Waltham, MA, USA) and Alexa-594 conjugated donkey anti-rabbit (1:250; Invitrogen, Waltham, MA, USA), respectively. In a separate set of experiments to validate the use of calbindin staining to identify PC numbers, immunohistochemistry was performed as described above for calbindin, followed by a counterstaining step using fluorescent Nissl stain NeuroTrace 435/455 (1:100; Life Technologies, Burlington, ON, Canada) for 1 hr. At least three slices per lobule were selected for each animal. Coronal slices were selected visually using images between 115 and 117 for anterior lobules, corresponding to lobules III, and images between 127 and 130 for posterior lobules, corresponding to lobules VIII/IX, according to the coronal view of the interactive Allen Mouse Brain Atlas viewer (Allen Institute of Brain Science, mouse coronal Atlas, 2011). Slice selection was further confirmed using cerebellar anatomical markers to ensure that data were compared from the same brain locations. This selection ensures that Purkinje cell density is quantified across the entire lobule, as the selected slices encompass the entire lobule.

### Cerebellar nuclei

Cerebellar sagittal slices were stained with primary antibodies: goat anti-Aldolase C (1:500, N-14; Santa Cruz Biotechnology, Dallas, TX, USA), rabbit anti-calbindin (1:1000, CB38A; Swant, Marly, Switzerland), and guinea pig anti-NeuN (1:500, ABN90; Millipore, Burlington, MA, USA). Primary antibodies were then tagged to secondary antibodies: Alexa-488 conjugated donkey anti-goat (1:250; Invitrogen, Waltham, MA, USA), Alexa-594 conjugated donkey anti-rabbit (1:250; Invitrogen, Waltham, MA, USA) and Alexa-647 conjugated donkey anti-guinea pig (1:250; Jackson Immuno, West Grove, PA, USA), respectively. For CN morphology analyses, the same sagittal sections described here were used. In a separate set of experiments to validate the use of calbindin staining as a proxy for PC innervation onto CN cells, immunohistochemistry was performed as described above for calbindin and NeuN, and additionally stained with primary antibody mouse anti-VGAT (1:500; Synaptic Systems, Göttingen, Germany), then tagged to secondary antibody Alexa-488 conjugated goat anti-mouse (1:250; Invitrogen, Waltham, MA, USA) to label GABAergic terminals in the CN. At least three slices per lobule were selected for each animal. Sagittal slices were selected visually using cerebellar anatomical markers to locate the fastigial and interposed nuclei, ensuring the same depth is chosen for comparisons.

### Confocal imaging and quantification

#### Cerebellar cortex

Images were acquired using a Zeiss LSM800 confocal fluorescent microscope equipped with Zen blue software (Zeiss, Oberkochen, Germany) using a 10x air objective and z-stacked for the depth of antibody penetration, consistent for all images. Images were tiled using Imaris stitcher software (Bitplane, Oxford Instruments) for whole lobule visualization. Microscope settings were kept constant during the imaging process for all conditions. Purkinje cell quantification was done on the projection image in IMARIS software, which combines the imaged z-stacks, ensuring quantification in the same depth and thickness of image. All visible Purkinje cells were counted. Calbindin labelling was used to count the total number of PCs. Zebrin labelling was used to distinguish between zebrin-negative (only positive for calbindin) and zebrin-positive puncta (positive for both calbindin and zebrin). Furthermore, zebrin labelling alone was used to calculate zebrin bandwidths by tracing a line parallel to the length of the lobule, midway of the length of the dendritic tree, as previously described (Toscano Márquez et al., 2021). For measurements of PC density, PC number was reported as a function of the PC layer length, reporting the number of cells per 100 µm, as previously described to account for differences in cerebellar volume(Toscano Márquez et al., 2021). Representative images are shown in pseudo-color.

#### Cerebellar nuclei

Images were acquired using a Zeiss LSM800 confocal fluorescent microscope equipped with Zen blue software (Zeiss, Oberkochen, Germany) using a 40x oil objective, and z-stacked for the depth of antibody penetration, consistent for all images. A digital zoom of 3x was used for analyses of PC innervation onto large CN cells using the projection image in IMARIS software, ensuring quantification in the same depth and thickness of image. Sampling the entire CN, 8-12 images per slice per region (anterior/posterior CN) were taken to ensure a representative quantification. Quantification of PC input onto CN cells was performed by counting individual calbindin-positive puncta of at least 1 µm in diameter, that were less than 0.5 µm from the NeuN-stained large CN cell (>12 µm diameter), where all visible puncta were counted. Calbindin puncta located more than 0.5 µm away from the NeuN-positive region were not included in the analysis as they were determined to not be forming a synapse with this CN cell. Colocalization of calbindin with zebrin was used to identify zebrin-positive PC innervation in the CN. Zebrin-negative innervation counts were obtained by subtracting the number of zebrin-positive innervation counts from the total number of innervation puncta. Sagittal slices from WT and CS mice used for cerebellar nuclei analyses were also used for CN morphology analyses, selecting only the NeuN channel for the latter. For CN morphology analyses, CN area and soma count were measured manually using ImageJ, where all visible cells were counted. Representative images are shown in pseudo-color.

#### Statistics

All imaging was acquired and analyzed while blinded to condition. Comparisons were made using Igor Pro 8.0 software. Data was first tested for normality, then analyzed using Student’s t-test for normally distributed data, or Mann-Whitney U test for non-normally distributed data. Data are represented by box and whisker plots, showing the median (horizontal line within boxes), 25^th^ and 75^th^ percentiles (rectangles) ± 1 SD (whiskers). Significance was determined as *p<0.05, **p<0.01, ***p<0.001, and p>0.05 if no comparison is shown.

## 3. RESULTS

### 3.1 Zebrin bands, PC density and CN innervation have formed normally in P25 CS mice

Previous studies reported a decrease in PC density in the anterior cerebellum of CS mice at age P57-P60 (Strømme et al., 2011; Xu et al., 2017), with zebrin-negative PCs being most susceptible (Strømme et al., 2011). To verify whether there are developmental changes, we set out to determine whether zebrin patterning develops normally in CS mice. We began our analyses at age P25, where mice are considered young adults (Strømme et al., 2011; Xu et al., 2017) when zebrin molecular differentiation of PCs is well-established (Hawkes & Herrup, 1995). As cerebellar degeneration in CS shows a selective pattern (Strømme et al., 2011; Xu et al., 2017), we compared two different cerebellar regions, anterior lobule III and posterior lobule VIII/IX, both of which are thought to be predominantly motor-associated lobules (Lin et al., 2020; Stoodley, Valera, & Schmahmann, 2012; Toscano Márquez et al., 2021). We stained coronal cerebellar slices from WT and CS mice with PC marker calbindin and zebrin to visualize the parasagittal compartmentalization of the cerebellum.

We quantified total PC density, as well as zebrin-negative and zebrin-positive PC densities, in both cerebellar regions at an age younger than 1 month old, an age when cerebellar area was unchanged between WT and CS mice (Xu et al., 2017). At age P25, we observed no significant differences in PC densities across the cerebellar cortex when comparing WT and CS mice (Student’s t-test for all comparisons other than anterior and posterior zebrin-positive densities, where Mann Whitney U tests were performed; anterior total density: p=0.5062; anterior zebrin-negative density: p=0.4036; anterior zebrin-positive density: p=0.708 (**Fig 1A-B**); posterior total density: p=0.8995; posterior zebrin-negative density: p=0.7435; posterior zebrin-positive density: p=0.996; **Fig 2A-B**), suggesting that PCs develop normally in CS mice.

**FIGURE 1.**
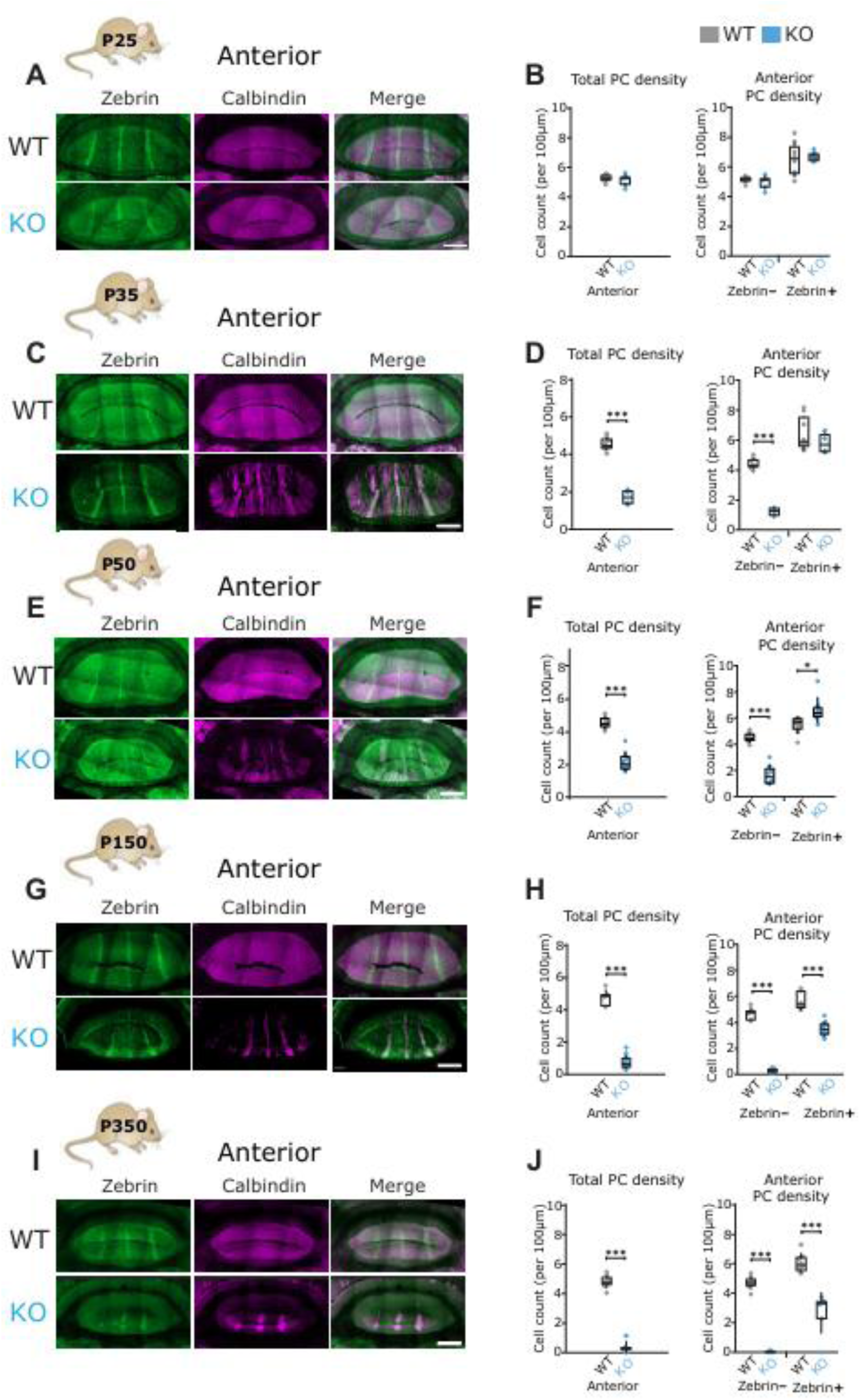
In the anterior, Purkinje cell death, limited to zebrin-negative Purkinje cells, begins at p35 and worsens progressively, ultimately affecting zebrin-positive cells at p150 in CS mice. (A, C, E, G, I) *Representative images of anterior lobule III in P25 (A), P35 (C), P50 (E), P150 (G), and P350 (I), from WT (top rows) and CS (bottom rows) mice, with PCs labelled with calbindin (purple) and zebrin (green). (B, D, F, H, J) Total PC density (left) and anterior lobule PC density (right) were calculated. N=4 for WT, N=3 for CS, 2 sections per lobule per animal (B, J); N=4 for WT, N=4 for CS, 2 sections per lobule per animal (D, F); N=3 for WT, N=5 for CS, 2 sections per lobule per animal (H). For statistics, student’s t-test for normally distributed data and Mann-Whitney U test for non-normally distributed data were used; *P<0.05, **P<0.01, ***P<0.001; n.s. refers to P>0.05. Scale bar (A, C, E, G, I), 400 µm*.

**FIGURE 2.**
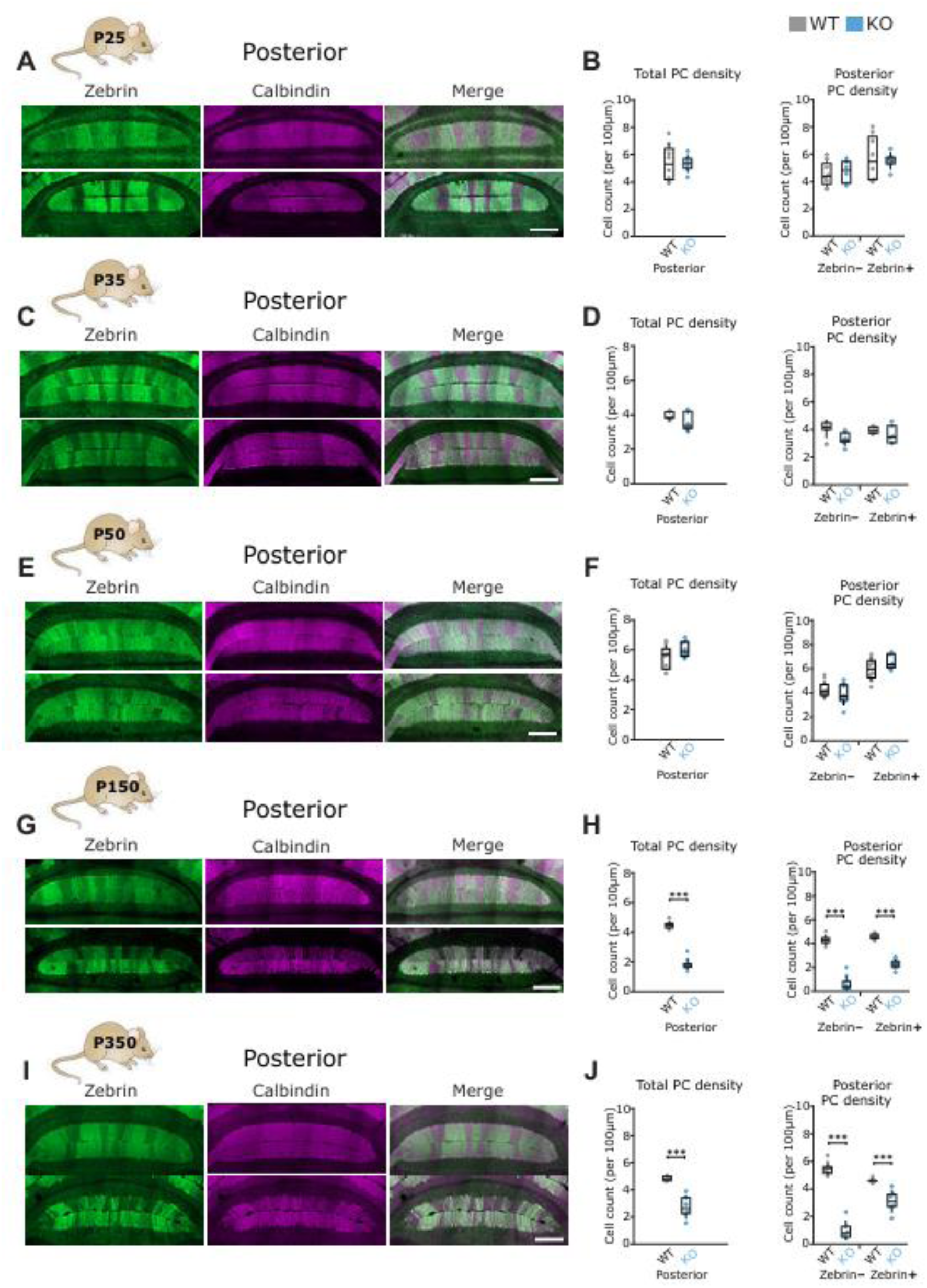
Posterior Purkinje cells are resilient to cell death until late disease stage, where both zebrin-negative and zebrin-positive cells are vulnerable to cell death starting at P150 in CS mice. (A, C, E, G, I) *Representative images of anterior lobule VIII in P25 (A), P35 (C), P50 (E), P150 (G), and P350 (I), from WT (top rows) and CS (bottom rows) mice, with PCs labelled with calbindin (purple) and zebrin (green). (B, D, F, H, J) Total PC density (left) and posterior lobule PC density (right) were calculated. N=4 for WT, N=3 for CS, 2 sections per lobule per animal (B, J); N=4 for WT, N=4 for CS, 2 sections per lobule per animal (D, F); N=3 for WT, N=5 for CS, 2 sections per lobule per animal (H). Scale bar (A, C, E, G, I), 300 µm*.

Next, we quantified zebrin-negative and zebrin-positive bandwidths (Refer to **Supplementary** Fig 1 f*or illustration of analysis*) and observed no significant differences in total zebrin-negative and zebrin-positive bandwidths, neither in the anterior (Student’s t-test; zebrin-negative bands: p=0.1023; zebrin-positive bands: p=0.1931; **Supplementary Fig 2A**) nor the posterior (Mann Whitney U test; zebrin-negative bands: p=0.379; zebrin-positive bands: p=0.368; **Supplementary Fig 2B**), when comparing WT and CS mice.

The CN are the final integration center of cerebellar information, receiving strong inhibitory input from PC axons, meaning that altered innervation onto the CN can strongly impact cerebellar function and motor control (Goodlett & Mittleman, 2017). Thus, we next sought to characterize PC innervation in the CN of CS mice. As the large projection neurons of the fastigial and interposed nuclei mainly receive input from the motor-associated vermal and paravermal regions, respectively (Amore et al., 2021; Apps & Hawkes, 2009; Voogd & Ruigrok, 2004), our analyses were based in these nuclei (Refer to **Supplementary** Fig 3 for illustration of CN innervation analysis). The pattern of zebrin projections is known to respect the anterior-posterior division of the CN, with the anterior CN region receiving predominantly zebrin-negative input and the posterior CN region receiving predominantly zebrin-positive input (Hawkes & Leclerc, 1986; Sugihara, 2011; Toscano Márquez et al., 2021), which we also observed (**Supplementary** Fig 3). Thus, we stained sagittal cerebellar vermal slices with NeuN to label the large cells of the CN, as well as the PC marker calbindin and zebrin, calculated the number of calbindin puncta in close contact with large CN cells, and subsequently quantified the zebrin identity of each PC putative synaptic contact in both anterior (**Fig 3C**) and posterior CN regions (**Fig 4C**).

**FIGURE 3.**
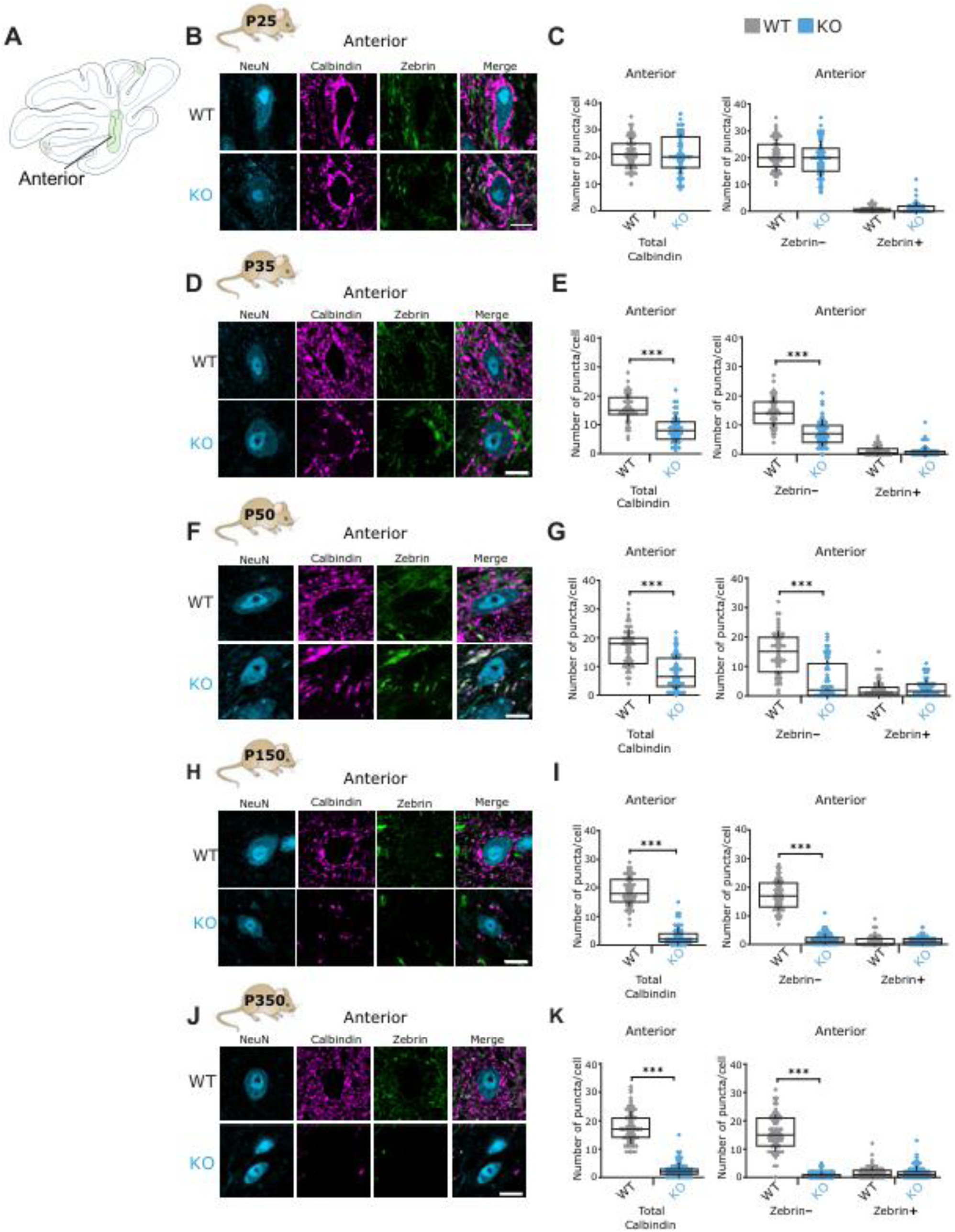
Inputs to the anterior CN follow a similar pattern to Purkinje cell degeneration, with no changes at P25, followed by a significant reduction in zebrin-negative inputs starting at P35. *A. Schematic representation of a sagittal cerebellar section showing the CN in the white matter with the anterior and posterior divisions. (B, D, F, H, J) Representative images of large CN neurons from the anterior CN region, innervated by PCs positive for calbindin (purple) and/or zebrin (green), from P25 (B), P35 (D), P50 (F), P150 (H) and P350 (J) WT (top rows) and CS (bottom rows) mice. (C, E, G, I, K) Total (left) and zebrin-specific (right) PC innervation onto large CN neurons in the anterior CN region were calculated. N=3 for WT, N=3 for CS, 2 sections per lobule per animal (C, E, G, I); N=4 for WT, N=4 for CS, 2 sections per lobule per animal (K). For statistics, student’s t-test for normally distributed data and Mann-Whitney U test for non-normally distributed data were used; *P<0.05, **P<0.01, ***P<0.001; n.s. refers to P>0.05. Scale bar (B, D, F, H, J), 10 µm*.

**FIGURE 4.**
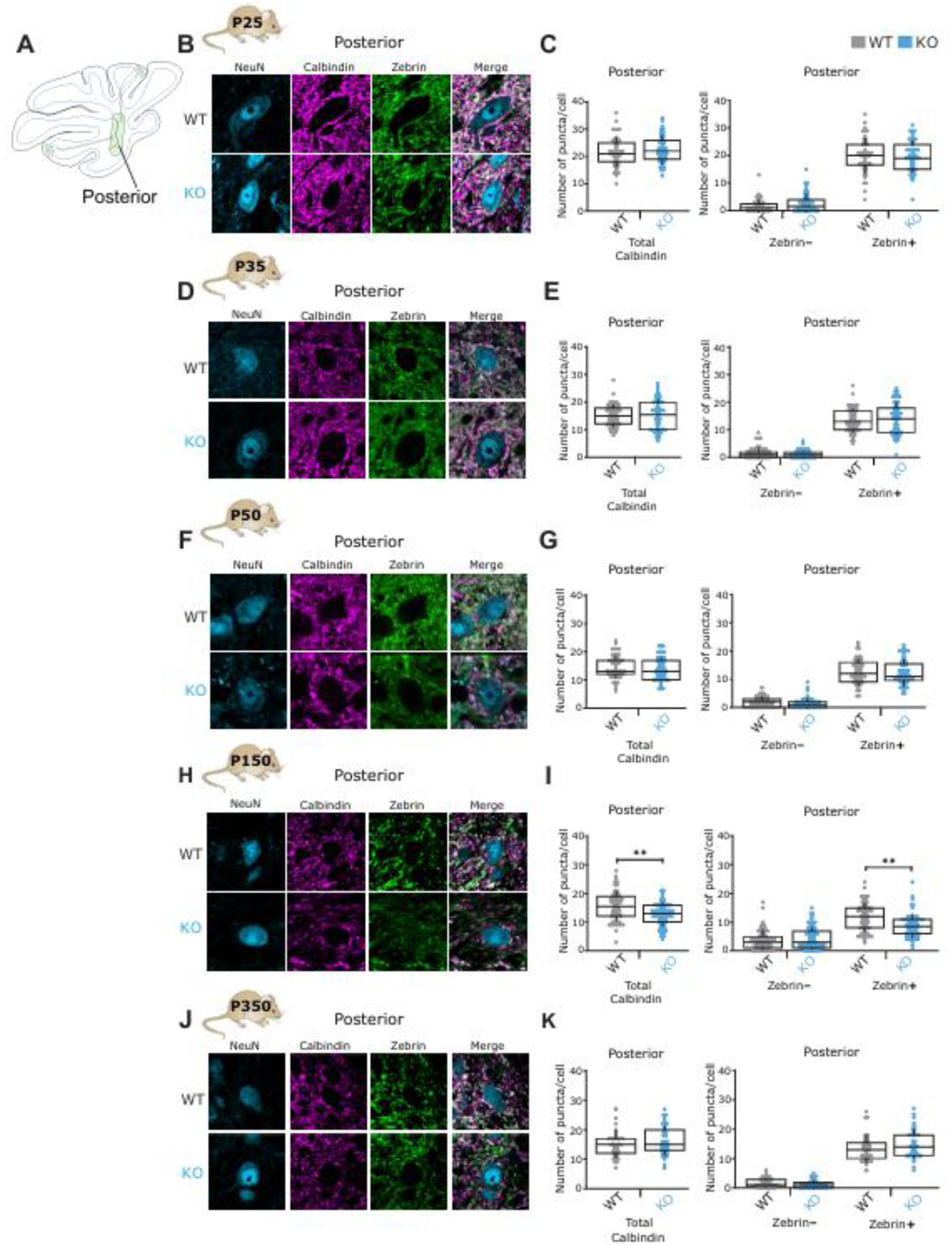
Purkinje cell inputs to the posterior CN are minimally affected, with only a transient decrease observed at p150 in CS mice. *A. Schematic representation of a sagittal cerebellar section showing the CN in the white matter with the anterior and posterior divisions. (B, D, F, H, J) Representative images of large CN neurons from the posterior CN region, innervated by PCs positive for calbindin (purple) and/or zebrin (green), from P25 (B), P35 (D), P50 (F), P150 (H) and P350 (J) WT (top rows) and CS (bottom rows) mice. (C, E, G, I, K) Total (left) and zebrin-specific (right) PC innervation onto large CN neurons in the posterior CN region were calculated. N=3 for WT, N=3 for CS, 2 sections per lobule per animal (C, E, G, I); N=4 for WT, N=4 for CS, 2 sections per lobule per animal (K). For statistics, student’s t-test for normally distributed data and Mann-Whitney U test for non-normally distributed data were used; *P<0.05, **P<0.01, ***P<0.001; n.s. refers to P>0.05. Scale bar (B, D, F, H, J), 10 µm*.

We observed no difference in PC innervation at P25 in either region of the CN in WT and CS mice (Student’s t-test for all comparisons other than anterior zebrin-positive innervation, where Mann Whitney U test was performed; anterior total input: p=0.7915; anterior zebrin-negative input: p=0.3063; anterior zebrin-positive input: p=0.071 (**Fig 3B-C**); Student’s t-test for all comparisons other than posterior zebrin-negative innervation, where Mann Whitney U test was performed; posterior total input: p=0.4543; posterior zebrin-negative input: p=0.1471; posterior zebrin-positive input: p=0.835; **Fig 4B-C**).

To ensure that calbindin is a good marker for functional PC terminals in the CN, we stained sagittal cerebellar vermal slices with the PC marker calbindin, as well as vesicular GABA transporter (VGAT) to label GABAergic terminals (Chaudhry et al., 1998). We observed no difference in the ratio of calbindin/VGAT puncta innervating large CN cells when comparing WT and CS mice (Student’s t-test; VGAT: p=0.719; **Supplementary** Fig 4).

### 3.2 Zebrin-negative PCs in the anterior cerebellum are more vulnerable to cell death in early disease progression (Ages P35 and P50)

Since the cerebellum of CS mice appears normal at age P25, we next determined whether zebrin patterning and PC density are altered ten days later, at age P35, as previous reports suggest that some PCs in the anterior cerebellum of some CS mice show markers of microglia/macrophages clustering around 1 month of age, which could be suggestive of neurodegeneration (Sikora et al., 2016). At this age, we observed a striking degeneration of anterior lobe zebrin-negative PCs (Student’s t-test for all comparisons other than anterior and posterior total PC density and posterior zebrin-negative PC density, where Mann Whitney U test was performed; anterior total density: p<0.0001; anterior zebrin-negative density: p<0.0001; anterior zebrin-positive density: p=0.2623 (**Fig 1C-D**); posterior total density: p=0.2869; posterior zebrin-negative density: p=0.1021; posterior zebrin-positive density: p=0.342; **Fig 2C-D**). In agreement with this data, we also observed a significant decrease in zebrin-negative bandwidth in the anterior cerebella of CS mice at this age (Mann Whitney U test; anterior zebrin-negative bands: p=0.003; anterior zebrin-positive bands: p=0.6093; **Supplementary Fig 2C**). The posterior cerebella of CS mice were largely spared at this age, aside from decreased bandwidth of zebrin-negative bands 2 (Student’s t-test; posterior zebrin-negative band 1: p=0.6298; posterior zebrin-negative band 2: p=0.0164; posterior zebrin-positive band 1: p=0.2074; posterior zebrin-positive band 2: p=0.435; posterior zebrin-positive band 3: p=0.614; **Supplementary Fig 2D**). These findings suggest a narrow window between P25 and P35 as the onset of anterior PC death in CS.

In the CN, we observed a significant decrease in total PC input in the anterior region of CS mice compared to WT at P35, affecting only zebrin-negative PC inputs (Student’s t-test for all comparisons other than anterior zebrin-positive innervation, where Mann Whitney U test was performed; anterior total input: p<0.0001; anterior zebrin-negative input: p<0.0001; anterior zebrin-positive input: p=0.645; **Fig 3D-E**). We observed no changes in PC innervation in the posterior CN region at this age between WT and CS mice (Mann Whitney U test for all comparisons; P35 posterior total input: p=0.966; P35 posterior zebrin-negative input: p=0.1344; P35 posterior zebrin-positive input: p=0.5661; **Fig 5D-E**).

**FIGURE 5.**
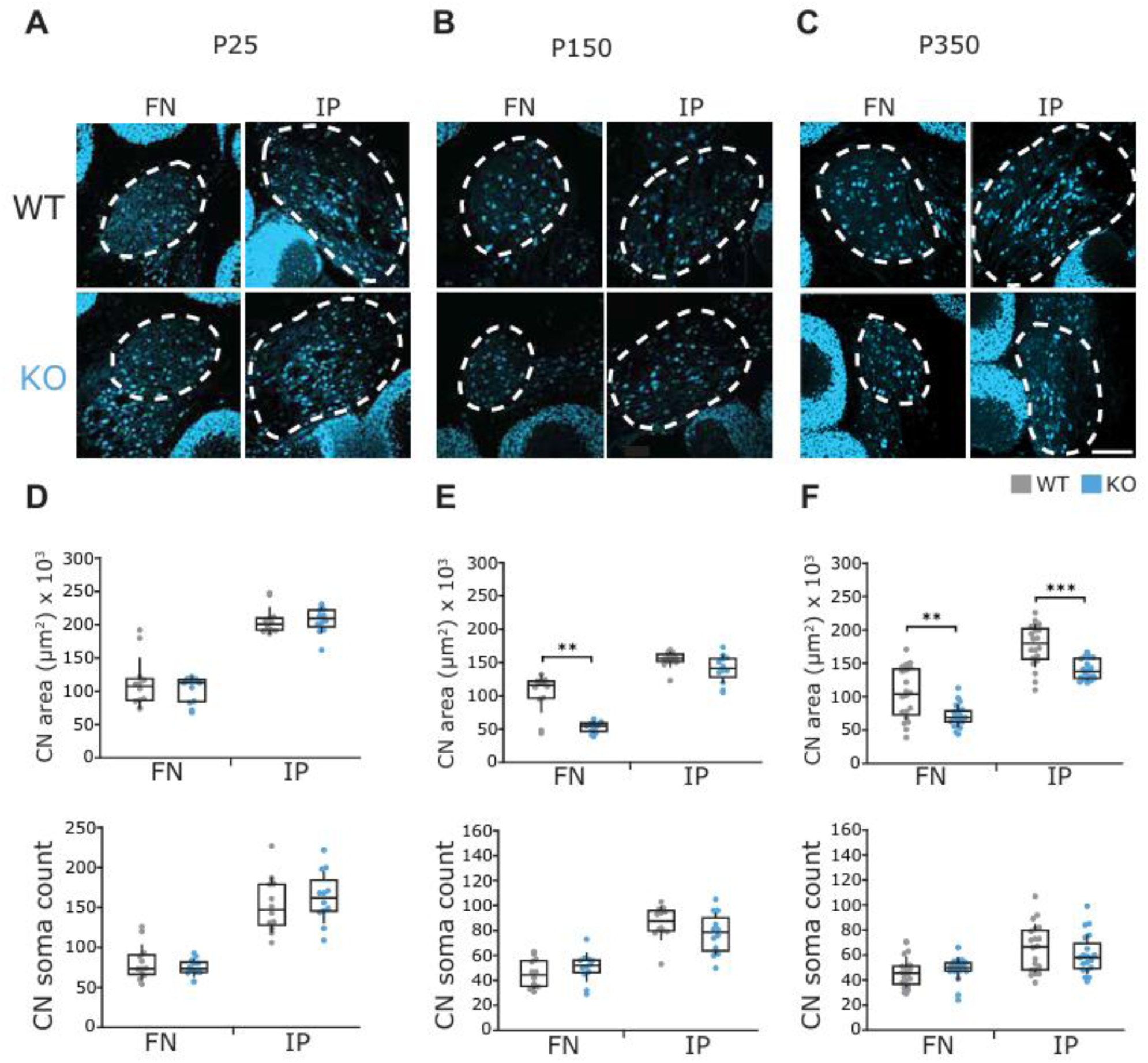
CN morphology alterations are apparent at age P150. (A-C) Representative images of fastigial (left) and interposed (right) CN nuclei of WT (top) and CS (bottom) mice at age P25 **(A)**, P150 **(B)**, and P350 **(C)**, labelled with NeuN (blue). **(D)** Area of the CN (top) and CN soma count (bottom) were calculated at age P25. No significant differences in area nor soma count were observed at this age, in both fastigial and interposed nuclei, when comparing CS (blue) and WT (grey) mice. **(E)** Area of the CN (top) and CN soma count (bottom) were calculated at age P150. At this age, the only significant change observed was a reduction in fastigial nuclei area, with no change in interposed nuclei area (top). No changes in CN soma count were observed between the two groups at this age (bottom). **(F)** Area of the CN (top) and CN soma count (bottom) were calculated at age P350. At this age, significant changes were observed in both fastigial and interposed nuclei area (top). No changes in CN soma count were observed between the two groups at this age (bottom). N=3 animals for WT, N=3 animals for CS, three sections per animal **(A, D)**; N=3 animals for WT, N=3 animals for CS, three sections per animal **(B, E)**; N=4 animals for WT, N=4 animals for CS, three sections per animal **(C, F)**. For statistics, Student’s t-test for normally distributed data and Mann Whitney U test for non-normally distributed data were used; *P<0.05, **P<0.01, ***P<0.001, P>0.05 if no comparison is shown. Scale bar, 150 µm **(C)**.

Next, we quantified PC density and zebrin bandwidths at age P50, an age prior to the previously reported onset of PC death in CS mice (Strømme et al., 2011; Xu et al., 2017). Here, we observed a significant decrease in anterior lobe zebrin-negative PC density of CS mice (Student’s t-test; anterior total density: p<0.0001; anterior zebrin-negative density: p<0.0001; **Fig 1E-F**). Interestingly, we also observed a significant increase in anterior lobe zebrin-positive PC density of CS mice (Student’s t-test; anterior zebrin-positive density: p=0.0177; **Fig 1E-F**). No changes in posterior lobe PC density were observed between WT and CS mice at this age (Student’s t-test; posterior total density: p=0.147; posterior zebrin-negative density: p=0.348; posterior zebrin-positive density: p=0.1404; **Fig 2E-F**). As seen at age P35, we observed a significant decrease in anterior lobe zebrin-negative bandwidth at P50 (Student’s t-test; anterior zebrin-negative bands: p=0.0004; **Supplementary Fig 2E**). We also observed a significant increase in anterior lobe zebrin-positive bandwidth in CS mice at this age, possibly further evidence of a compensatory effect in CS cerebella at this age (Student’s t-test; anterior zebrin-positive bands: p=0.0072; **Supplementary Fig 2E**). Posterior lobe bandwidths were largely unchanged at age P50 in CS mice, aside from a significant decrease in zebrin-negative bands 2 (Student’s t-test for all comparisons other than posterior zebrin-negative band 1 and posterior zebrin-positive band 2, where Mann Whitney U test was performed; posterior zebrin-negative band 1: p=0.3624; posterior zebrin-negative band 2: p=0.0058; posterior zebrin-positive band 1: p=0.9598; posterior zebrin-positive band 2: p=0.064; posterior zebrin-positive band 3: p=0.6104; **Supplementary Fig 2F**).

Since we used PC marker calbindin for our analyses, any decreases observed could be deemed simply a downregulation of the calcium-binding protein itself. Thus, to test our hypothesis that the loss of calbindin was indeed due to cell death, we stained P50 coronal cerebellar slices with calbindin as well as a Nissl stain, which is commonly used as a general label for neuronal nuclei. In doing so, we were able to confirm that the loss of calbindin reflected a loss of PCs, and not simply a decreased calbindin signal (Student’s t-test for all comparisons; WT: p=0.757; CS: p=0.272**; Supplementary** Fig 5).

Alterations in the anterior CN region of CS mice continued at age P50, where we observed a similar decrease in zebrin-negative PC innervation, with no change in zebrin-positive inputs (Mann Whitney U test; anterior total input: p<0.0001; anterior zebrin-negative input: p<0.0001; anterior zebrin-positive input: p=0.169; **Fig 3F-G**). No changes were observed in the posterior CN PC innervation between WT and CS mice at this age (Mann Whitney U test for all comparisons other than P50 posterior total input and P50 posterior zebrin-negative input, where Student’s t-test was performed; P50 posterior total input: p=0.367; P50 posterior zebrin-negative input: p=0.278; P50 posterior zebrin-positive input: p=0.064; **Fig 4F-G**).

### 3.3 PC death extends to the posterior cerebellum in later disease stages, where both zebrin-negative and zebrin-positive PCs are equally affected (Ages P150 and P350)

Though studies on patterned PC vulnerability have highlighted PC resiliency in the posterior cerebella of CS mice, it remains unclear how zebrin identity influences posterior PC viability in CS mice and what the timeline of this resilience is. To address this, we analyzed PC density and zebrin bandwidths from age P150 WT and CS mice, an age previously linked to posterior degeneration in CS (Strømme et al., 2011). We observed anterior lobe zebrin-positive PC degeneration in CS mice in later disease progression, with significant decreases in both zebrin-negative and zebrin-positive anterior lobule PC densities (Mann Whitney U test; anterior total density: p<0.0001; anterior zebrin-negative density: p<0.0001; anterior zebrin-positive density: p<0.0001; **Fig 1G-H**). Interestingly, we report that the posterior cerebellar degeneration in CS mice present at age P150 affects both zebrin-negative and zebrin-positive PCs (Mann Whitney U test; posterior total density: p<0.0001; posterior zebrin-negative density: p<0.0001; posterior zebrin-positive density: p<0.0001; **Fig 2G-H**). As observed at earlier timepoints, we report a decreased zebrin-negative bandwidth in the anterior cerebella of CS mice at age P150, without changes in anterior lobe zebrin-positive bandwidth (Mann Whitney U test for anterior zebrin-negative bands, Student’s t-test for anterior zebrin-positive bands; anterior zebrin-negative bands: p<0.0001; anterior zebrin-positive bands: p=0.069; **Supplementary Fig 2G**). In the posterior, we observed alterations in both zebrin-negative and zebrin-positive bandwidths; with decreased bandwidth of zebrin-negative bands 2 and zebrin-positive band 1 (Mann Whitney U test for all comparisons other than posterior zebrin-positive bands 2 and 3, where Student’s t-test was performed; posterior zebrin-negative band 1: p=0.288; posterior zebrin-negative bands 2: p=0.0016; posterior zebrin-positive band 1: p=0.0026; posterior zebrin-positive bands 2: p=0.8045; posterior zebrin-positive bands 3: p=0.479; **Supplementary Fig 2H**).

Next, we analyzed PC innervation in the CN at age P150. Here, we report a significant decrease in PC innervation in the anterior CN region of CS mice, affecting only zebrin-negative PC inputs (Mann Whitney U test for all comparisons; P150 anterior total input: p<0.0001; P150 anterior zebrin-negative input: p<0.0001; P150 anterior zebrin-positive input: p=0.302; **Fig 3H-I**). Interestingly, despite observing decreases in both zebrin-negative and zebrin-positive PC densities in the cerebellar cortex of CS mice at age P150, decreased PC innervation in the posterior CN region of CS mice at age P150 was restricted to zebrin-positive PC inputs only, with no changes in zebrin-negative PC inputs (Mann Whitney U test for all comparisons; posterior total input: p=0.0024; posterior zebrin-negative input: p=0.507; posterior zebrin-positive input: p=0.0011; **Fig 4H-I**).

Finally, we moved to a very late timepoint in disease progression, age P350, where lobule IX degeneration was previously reported in CS mice (Xu et al., 2017). We observed decreased PC density in CS mice affecting both zebrin-negative and zebrin-positive PCs in anterior and posterior lobules (Mann Whitney U test for all comparisons other than anterior total PC density; anterior total density: p<0.0001; anterior zebrin-negative density: p<0.0001; anterior zebrin-positive density: p=0.0006 (**Fig 1I-J**); posterior total density: p<0.0001; posterior zebrin-negative density: p<0.0001; posterior zebrin-positive density: p=0.0009; **Fig 2I-J**). Additionally, we observed a decreased zebrin-negative bandwidth and an increased zebrin-positive bandwidth in the anterior cerebella of CS mice (Mann Whitney U test for anterior zebrin-negative bands, Student’s t-test for anterior zebrin-positive bands; anterior zebrin-negative bands: p=0.0008; anterior zebrin-positive bands: p<0.0001; **Supplementary Fig 2I**). Despite reporting considerable PC loss, we observed no changes in zebrin bandwidths in the posterior cerebella of CS mice at age P350 (Mann Whitney U test for all comparisons; posterior zebrin-negative band 1: p=0.5363; posterior zebrin-negative bands 2: p=0.644; posterior zebrin-positive band 1: p=0.758; posterior zebrin-positive bands 2: p=0.968; posterior zebrin-positive bands 3: p=0.0522; **Supplementary Fig 2J**).

Analyzing CN innervation at age P350, we observed a significant decrease in PC innervation of the anterior CN region in CS mice, affecting only zebrin-negative PC inputs (Mann Whitney U test for all comparisons; P350 anterior total input: p<0.0001; P350 anterior zebrin-negative input: p<0.0001; P350 anterior zebrin-positive input: p=0.913; **Fig 4J-K**). We observed no changes in PC innervation in the posterior CN region at age P350 when comparing WT and CS mice, despite observing extensive degeneration in the posterior cerebellar cortex of CS mice at this age (Student’s t-test for all comparisons; posterior total input: p=0.3593; posterior zebrin-negative input: p=0.0922; posterior zebrin-posterior input: p=0.145; **Fig 5J-K**).

### 3.4 No death of CN neurons in the fastigial and interposed nuclei is observed in CS

As PC innervation makes up most of the synaptic contact on CN neurons (Uusisaari & Knöpfel, 2011), we hypothesized that the considerable cerebellar degeneration in the cerebellar cortical vermis of CS mice would cause large-scale morphological changes in the CN, with a loss of large CN neurons in later disease progression. To investigate this, we used the neuronal marker NeuN to determine CN area and CN soma count, comparing WT and CS mice at different ages. As expected, we did not find differences in CN area nor cell number at P25, consistent with the idea that the cerebellum develops normally in CS (Student’s t-Test for all comparisons; FN area: p=0.9661; IP area: p=0.4863; FN soma count: p=0.3498; IP soma count: p=0.4891; **Fig 5A, D)**. Consistent with cerebellar atrophy seen in patients and mice (Pescosolido et al., 2014; Xu et al., 2017), we found reduced fastigial nucleus area at P150 (Student’s t-Test for all comparisons; FN area: p=0.0001; IP area: p=0.0704; FN soma count: p=0.2724; IP soma count: p=0.1824; **Fig 5B, E)** and reduced fastigial and interposed areas at P350 (Student’s t-Test for all comparisons except NF soma count; FN area: p<0.0001; IP area: p=0.0002; FN soma count: p=0.2079; IP soma count: p=0.3736; **Fig 5C-F**). Surprisingly, although we observed a reduced CN area at ages P150 and P350, we did not observe any changes in CN soma count between WT and CS mice (Fig 5E, F). This data shows that despite the large loss of innervation, the CN neurons remain resilient.

### 3.5 PC spontaneous firing patterns develop normally in P25 CS mice, with deficits evident at age P35

In many mouse models of ataxia, alterations in PC spontaneous firing are seen prior to motor deficits and PC death, and often serve as the first sign of cerebellar dysfunction (Cook, Fields, & Watt, 2021). To address whether changes in PC spontaneous spiking rate contribute to early stages of cerebellar ataxia in CS mice, we performed cell-adjacent extracellular recordings from PCs to non-invasively measure their spontaneous firing patterns with the addition of blockers of fast synaptic transmission to ensure we record the intrinsic firing rates, unaffected by any inputs (Ady et al., 2018). To identify when alterations in PC firing patterns arise, we chose two timepoints prior to when others have reported PC loss in CS, which correspond to the timepoints when we first saw no cell death and when we first saw significant cell death: P25 and P35. We chose to record from two locations associated with motor control (Lin et al., 2020; Stoodley, Valera, & Schmahmann, 2012): anterior lobule III, a region that is known to exhibit PC cell death early, and posterior lobule VIII, a region that we found to be resilient to cell death at these ages (Xu et al., 2017). We measured two properties of PC firing that have been reported to be altered in several ataxias (Cook, Fields, & Watt, 2021): PC spontaneous firing frequency, as well as the coefficient of variation of the inter-spike interval (CV) of PC action potentials, a measure of PC firing regularity.

At a timepoint prior to cerebellar degeneration in CS (Strømme et al., 2011; Xu et al., 2017), P25, which is thought to reflect a largely mature cerebellar circuit (Arancillo et al., 2014; Kaneko et al., 2011) we observed no differences in the firing properties of PCs in CS and WT mice in either lobule and a similar distribution of firing frequencies (**Fig 5D-E**) (Student’s t-test for all comparisons other than lobule III firing frequency, where Mann Whitney U test was performed; III frequency: p=0.2283; VIII frequency: p=0.521; III regularity: p=0.618; VIII reg: p=0.7524; **Fig 6B**). Since we found that PC death in CS mice is significant at p35, we chose to look at this timepoint to see if firing deficits may occur around the same time. Thus, we next analyzed PC intrinsic firing patterns at age P35. At this timepoint, we observed a significant decrease in the firing frequency of lobule III PCs in CS mice with no changes in the firing regularity, where we observed a larger proportion of cells firing at frequencies less than 40Hz, with none firing at over 80Hz compared to WT where the majority of cells fired at frequencies over 40Hz, with about half of the cells firing over 60Hz (**Fig 6F-G**) (Student’s t-test for firing regularity, Mann Whitney U test for firing frequency; III frequency: p<0.0001; VIII frequency p=0.6504; III regularity: p=0.1991; VIII regularity: p=0.2757; **Fig 6C**).

**FIGURE 6.**
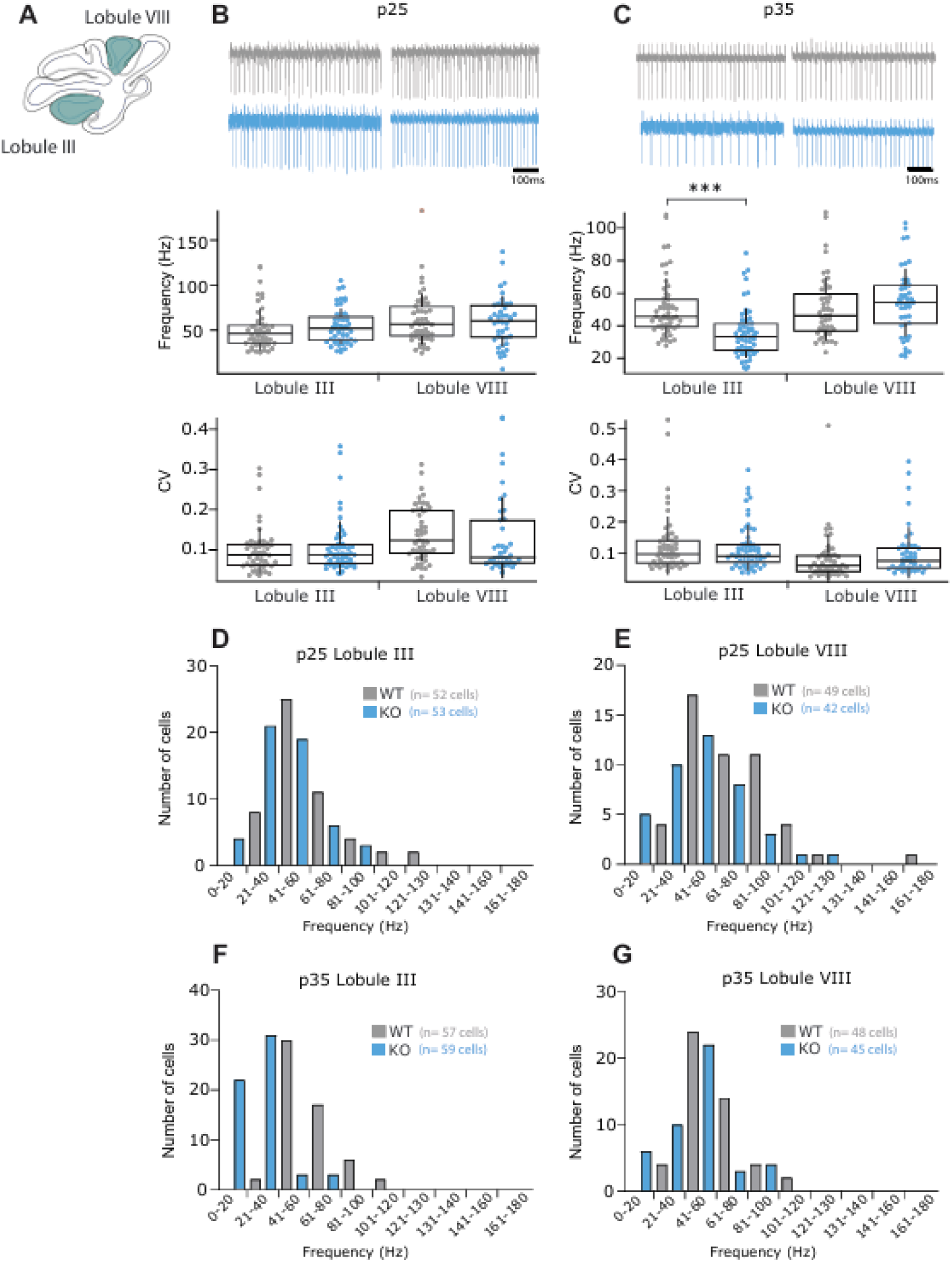
Intrinsic Purkinje cell firing deficits are observed at p35 only in the anterior vulnerable region. **(A)** Schematic representation showing anterior lobule III and posterior lobule VIII recording site. **(B-C)** Sample spike trains from a representative P25 **(B)** and P35 **(C)** WT (grey) and CS (blue) PC from anterior lobule III and posterior lobule VIII (top). The frequency (top graphs), reported as spikes per second (Hz), and regularity (bottom graphs), reported as coefficient of variation (CV), of PC firing were recorded and analyzed at ages P25 **(B)** and P35 **(C)**, in both anterior lobule III and posterior lobule VIII following the application of synaptic blockers. (**D-G).** Histograms demonstrating the distribution of firing frequencies of recorded PCs in the anterior lobule III at P25 **(D)** and P35 **(F)**, and in the posterior lobule VIII at P25 **(E)** and P35 **(G)**, showing a reduced number of high firing PCs at P35 in the anterior lobule only. (N=number of mice, n=number of cells) N=4, n=52 for WT; N=4, n=53 for CS **(B, lobule III)**. N=4, n=49 for WT; N=4, n=42 for CS **(B, lobule VIII)**. N=4, n=57 for WT; N=4, n=59 for CS **(C, lobule III)**. N=4, n=48 for WT; N=4, n=45 for CS **(C, lobule VIII)**. For statistics, student’s t-test for normally distributed data and Mann Whitney U test for non-normally distributed data were used; *P<0.05, **P<0.01, ***P<0.001; n.s. refers to P>0.05.

## 4. DISCUSSION

While most research efforts on CS have focused on brain regions such as the hippocampus (Gao et al., 2019; Xu et al., 2017), the ataxic pathophysiology of CS has been less studied. In this study, we show that the patterned vulnerability of PCs, which has been previously described in CS mice (Strømme et al., 2011; Xu et al., 2017), is driven by zebrin molecular identity and cerebellar location. While PCs of the anterior vermis expressing zebrin show greater resilience compared to their zebrin-negative counterparts, we provide evidence that these resilient PCs do eventually degenerate in CS, albeit at a slower pace (see Fig 7 and **Table 1** for summaries). Though zebrin identity dictates PC vulnerability in the anterior CS cerebella, both zebrin-negative and zebrin-positive PCs of the posterior vermis appear vulnerable to degeneration in later disease stages, independent of zebrin molecular identity. Importantly, while CS is thought to be a mixed neurodevelopmental and neurodegenerative condition (Xu et al., 2017), the development of cerebellar PC patterning and PC firing rates in young juvenile CS mice appears normal, suggesting that cerebellar development may be dissociated from disease pathophysiology. Projections from PCs to target neurons in the CN shared a similar pattern of degeneration with that seen in the cerebellar cortex, with reduced synaptic innervation observed only among zebrin-negative PC puncta in the anterior CN region early in disease progression, while both zebrin-negative and zebrin-positive PC puncta in the posterior CN region shared a greater resiliency in later disease stages. These findings highlight that both zebrin molecular identity and cerebellar location influence PC vulnerability in CS mice.

**FIGURE 7.**
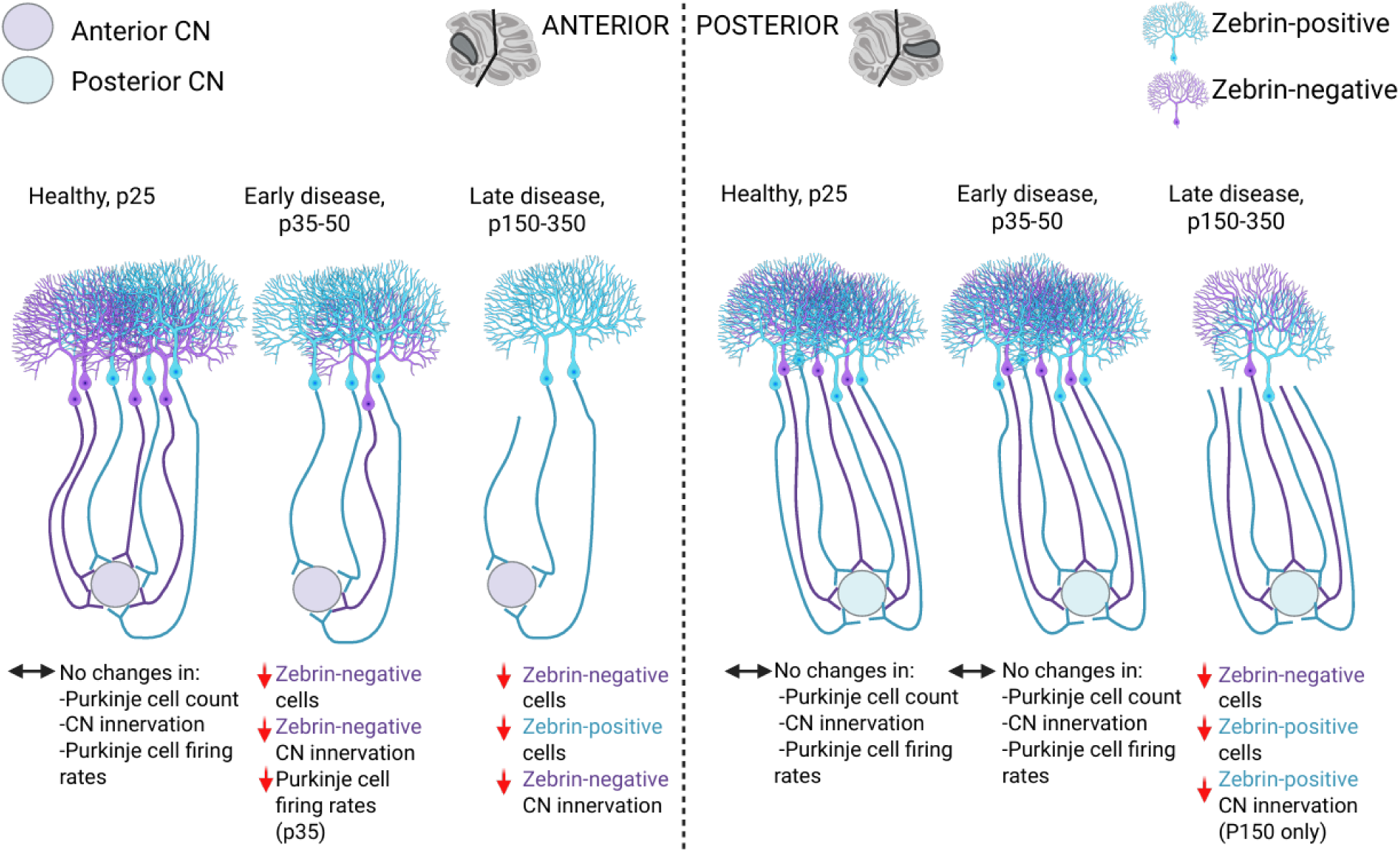
Summary schematic illustrating the findings related to Purkinje cell count, Purkinje cell innervation of the CN and Purkinje cell firing. ***The cerebellum appears to develop normally, with no loss in Purkinje cell or Purkinje cell innervations to the CN, and no changes in Purkinje cell firing rates at p25. Deficits start at p35, with the onset of zebrin-negative Purkinje cell death in the anterior only, accompanied by loss of zebrin-negative innervation to the anterior CN region, and the onset of firing deficits in the anterior vulnerable region, but not in the posterior resilient region. At late disease stages, starting at p150, Purkinje cell loss occurs both in the anterior and posterior cerebellum, independently of zebrin molecular identity. However, Purkinje cell innervation loss to the CN remains localized to the anterior CN and specific to zebrin-negative Purkinje cells*.**

**TABLE 1.** Summary table showing changes in Purkinje cell count, Purkinje cell innervation of the CN, and firing rates and regularity of Purkinje cells. *No changes in any parameters investigated are observed at P25. Deficits in PC firing rates commence at P35 only in the anterior cerebellum, around the same time as the death of zebrin-negative Purkinje cells occurs in that region. A possible transient upregulation of zebrin-positive cells in the anterior is observed at P50, followed by a loss of PCs independent of their molecular identity at P150, worsening at P350. Only zebrin-negative Purkinje cell innervations to the anterior CN are reduced throughout the disease progression, despite zebrin-positive cell loss later in the disease. The posterior Purkinje cells are spared until late disease stages, where death of Purkinje cells, independently of their molecular identity, is observed. PC innervations to the posterior CN are transiently reduced at p150, and not observed anymore at p350*.

|  | P25 |  | P35 |  | P50 |  | P150 |  | P350 |  |
| --- | --- | --- | --- | --- | --- | --- | --- | --- | --- | --- |
|  | Anterior | Posterior | Anterior | Posterior | Anterior | Posterior | Anterior | Posterior | Anterior | Posterior |
| **PC count** | No change | No change | Zebrin-↓ | No change | Zebrin-↓<br>Zebrin+↑ | No change | Zebrin-↓<br>Zebrin+↓ | Zebrin-↓<br>Zebrin+↓ | Zebrin-↓<br>Zebrin+↓ | Zebrin-↓<br>Zebrin+↓ |
| **CN innervation** | No change | No change | Zebrin-↓ | No change | Zebrin-↓ | No change | Zebrin-↓ | Zebrin+↓ | Zebrin-↓ | No change |
| **PC firing rates** | No change | No change | Decrease<br>↓ | No change |  |  |  |  |  |  |
| **Regularity of PC firing** | No change | No change | No change | No change |  |  |  |  |  |  |

One of the important findings from this study is the observation of a narrow and aggressive window of vulnerability for PC death in the cerebella of CS mice, occurring between the ages P25-P35. This 10-day window is highlighted not only by cerebellar cortex PC degeneration and loss of CN synaptic innervation, but also in the intrinsic firing patterns of PCs, with anterior lobe PCs of CS mice exhibiting decreased firing frequency at age P35. More studies will be needed to elucidate the potential mechanisms which may be triggering such a rapid and aggressive insult in the CS cerebellum. Interestingly, axonal spheroids were observed in Purkinje cells in CS mice as early as p19, before their degeneration, and before intrinsic firing deficits, as we have shown that they start at p35, but their role in affecting Purkinje cell vulnerability and firing deficits in CS is unknown (Strømme et al., 2011). Axonal spheroids of Purkinje cells, also called torpedoes, have been found both in healthy development and diseased states, such as in lysosomal storage diseases (Steven et al., 2010). In healthy states, they occur transiently during development, peaking at p11, and have been reported to increase the fidelity of axon potential propagation (Lang-Ouellette et al., 2021; Ljungberg et al., 2016). The firing rates of Purkinje cells were not different between Purkinje cells containing the swellings and those without them (Lang-Ouellette et al., 2021). Perhaps, their role in disease state may be different. In many lysosomal storage diseases, Purkinje cells were found to die following the formation of the swellings, and it was suggested that these swellings prevent proper retrograde transport in Purkinje cell axons, thereby making them more vulnerable to cell death (Steven et al., 2010). In a mouse model of spinocerebellar ataxia type 6 (SCA6), elevated torpedo numbers were seen in adult mice at the age of Purkinje cell loss and firing deficits, suggesting that torpedoes may be linked with aberrant firing rates (Grömmke et al., 2025; Ljungberg et al., 2016). However, in CS, the number of axonal swellings was not quantified and not compared to WT mice, making it unclear whether the number of swellings is significantly different. Axonal swellings, although peaking at p11, were still found in about 20% of Purkinje cells as late as p30 (Ljungberg et al., 2016), making it unclear whether the swellings seen at p19 in CS are pathological or not, as their contents have not been visualized by electron microscopy (dense packing of intracellular organelles is not seen in developmental swellings, but has been shown in disease (Lang-Ouellette et al., 2021)). In heterozygous females, axonal swellings were seen on the remaining Purkinje cells at 32 weeks (Sikora et al., 2016), which could suggest that the swellings do improve axonal propagation, and may help these neurons survive, which is consistent with previous work suggesting that Purkinje cells with axonal swellings are the surviving neurons, although they may also degenerate later (Louis et al., 2014). Since the axonal swellings do not affect the firing rates of Purkinje cells in healthy mice, we hypothesize that they also will not affect the firing rates of Purkinje cells in CS, but might instead attempt to improve action potential propagation to the CN, which may be neuroprotective, or may, on the contrary, be disease-causing and contribute to Purkinje cell vulnerability by preventing proper retrograde trafficking independently of firing rates. More research is needed to determine whether axon swellings are disease-causing or are neuroprotective in CS.

Patterned PC death has been observed in several different ataxias and neurological disorders caused by various genetic insults (Sarna & Hawkes, 2003) and even aging (Donofrio et al., 2025). In fact, one of the earliest reported and most explored defects is acid-sphingomyelinase deficiency (Niemann-Pick disease AB, Sarna et al. 2001) and NPC1 deficiency (Niemann-Pick disease type C, Sarna et al. 2003). Importantly, patterned PC degeneration in the latter model has been documented as cell-autonomous (ref papers from Scott and Maue labs). Although not presented in such detail, it is more than likely that the cell-autonomous pathogenesis applies to a broad range of conditions (CS included) characterized by the shared PC neurodegenerative pattern. It would be interesting to see if zebrin plays a role in this phenomenon.

While zebrin-negative PCs appear more vulnerable in CS (Strømme et al., 2011), a pattern which is shared by other ataxia models (Sarna & Hawkes, 2003), our study highlights that not all zebrin-negative PCs are equal, with those residing in the anterior vermis being particularly susceptible to death in CS. While this finding has also been noted in ARSACS (Toscano Márquez et al., 2021), zebrin molecular identity and its impact on patterned cerebellar degeneration remain largely underappreciated. The differential influence of zebrin molecular identity depending on cerebellar location may be due to additional molecules of cerebellar compartmentalization. Indeed, we now understand that heterogeneity in the cerebellum is more complex than previously appreciated (Apps et al., 2018; Cerminara et al., 2015; Wu et al., 2019). For example, flocculonodular lobule X, where PCs are uniformly zebrin-positive (Lin et al., 2020), has been described as a naturally resilient cerebellar region in several models of ataxia (Hernández-Pérez, Weruaga, & Díaz, 2023; Slemmer et al., 2007). In a separate set of experiments, we observed no difference in the PC density of lobule X between WT and CS mice at age P350 (**Supplementary** Fig 6), a finding previously reported at an earlier age in CS (Xu et al., 2017). In addition to being uniformly zebrin-positive, PCs of lobule X are also characterized by the expression of various molecular markers (Blot et al., 2021; Duffin et al., 2010; Wu et al., 2019), as well as unique circuitry motifs, with axons of lobule X PCs known to bypass the CN to synapse directly onto neurons in the vestibular nuclei (Sugihara, 2011). Together, these findings indicate that the patterned vulnerability of cerebellar PCs in CS likely extends beyond the identity of zebrin and will require further research attention.

At ages P35 and P50, we observed a significant decrease in the bandwidth of posterior lobule zebrin-negative band 2 in CS mice. This may be indicative of the posterior lobe attempting to compensate for deficits occurring in the anterior CS cerebellum at this age. More studies will be needed to elucidate the specific roles played by individual zebrin bands, and how these different parasagittal compartments interact to control motor behaviour. The ability of CS mice to walk at all at advanced disease stages (Strømme et al., 2011), combined with our finding that very few cells are left intact in the anterior lobe in the late stages of disease progression, suggests a strong compensatory event that may not be well explained by zebrin-positive compensation in the anterior lobe alone, considering the limited number of these PCs in the anterior vermis, as well as our finding that these cells also eventually undergo degeneration. In fact, this may be better explained by the activity of the posterior lobe, which becomes affected much later in disease progression. More specifically, compensation in the flocculonodular lobe should be further explored in CS cerebella, as this region has been shown to be particularly resilient in many cases of ataxia (Hernández-Pérez, Weruaga, & Díaz, 2023; Slemmer et al., 2007), possibly due to its uniform zebrin-positive PC expression and unique circuitry (Sugihara, 2011).

PCs are the only output of the cerebellar cortex, and innervate the large cells of the CN to communicate with other brain regions (Voogd, Jaarsma, & Marani, 1996). Any changes in PC innervation onto the CN will likely have devastating effects on motor control. Thus, we set out to characterize PC innervation in the CN in CS mice, as this has not been tested before. Importantly, as PC innervation in the CN is known to respect the anterior-posterior boundary set by zebrin expression (Hawkes & Leclerc, 1986; Sugihara, 2011; Toscano Márquez et al., 2021), we investigated both total PC input as well as zebrin-negative and zebrin-positive PC input in the CN. We found that in CS mice, alterations in CN innervation largely match our observations seen in the cerebellar cortex, with seemingly normal development of CN innervation in young adult mice at P25, and loss of zebrin-negative PC innervation in the anterior CN region starting at age P35, further highlighting the 10-day window of vulnerability in the anterior CS cerebella. Interestingly, we obtained conflicting findings in CN innervation at later ages. At age P150, we report decreased total PC innervation in the posterior CN region, with only changes in zebrin-positive PC inputs being observed, yet no alterations were detected at age P350. These conflicting findings may be explained by animal variation since coronal and sagittal slices for the different analyses originated from separate animals.

For the first time to our knowledge, CN neuronal counts were done in fastigial and interposed nuclei in a mouse model of CS. Interestingly, despite a significant loss of PC contact, no cell death was seen, showing that the large CN neurons remain highly resilient in CS. This is consistent with human data, where mild neuronal loss was reported only in the dentate nucleus (Garbern et al., 2010). Importantly, in the *shaker rat*, an ataxia model caused by a spontaneous mutation in the *SLC9A6* gene which shares many characteristics with CS (Figueroa et al., 2023), it was shown that deep brain stimulation (DBS) in the CN, downstream from degenerating PCs, can ameliorate cerebellar pathology and the ataxic phenotype (Anderson et al., 2019). Since DBS relies on the sufficient presence of remaining neurons for stimulation (Anderson et al., 2019), our findings of highly resilient large CN neurons in CS may open an avenue for therapeutic intervention using DBS in CS ataxia.

We focus this paper on only male mice, as we wanted to see a strong phenotype and characterize it. However, heterozygote females would be interesting and important to investigate as well. In the human carriers, a range of phenotypes were seen, ranging from no problems seen to a multitude of different characteristics, usually psychiatric in nature, where female carriers showed learning and behavioural issues, including mild to moderate intellectual disability, attention deficit/hyperactivity disorder, schizophrenia and autism (Christianson et al., 1999; Gilfillan et al., 2008; Pescosolido et al., 2014), although one of the female carriers presented with microcephaly, and another carrier had mild truncal ataxia (Pescosolido et al., 2014; Schroer et al., 2010). Informative previous work was done by Sikora et al. (2016) studying female CS heterozygotes in a mouse model, where mosaicism in β-Gal expression, replacing NHE6, was shown (Sikora et al., 2016). They showed less severe Purkinje cell degeneration compared to males, but in the same location and at the same time, with significant death in the anterior lobule

III but not the flocculonodular lobule X at 32 weeks of age. They also found neurono/dendritophagic clustering of CD68^+^ macrophages/microglia on some Purkinje cells at 4 weeks of age, suggesting some Purkinje cell death, but less in the heterozygotes compared to the males. However, this was not quantified and was not compared to WT, making it unclear whether significant Purkinje cell death starts later in heterozygotes compared to male CS mice. Considering that the severity is lowered in heterozygotes, we hypothesize that significant cell death would be seen a little later in heterozygotes, but it is a worthy focus in future studies.

## 6 ACKNOWLEDGMENTS

We thank all past and present members of the McKinney and Watt labs for constructive and thoughtful feedback on the project. Imaging was performed at the McGill University Advanced BioImaging Facility (ABIF), and we thank ABIF staff members for their technical support. We are grateful for the animal care and training we received from the McGill Animal Resources Centre (CMARC), and particularly for the expert help of Tanya Koch.

## Contributions

R.A.M., and A.J.W. designed the research; LCM., J.T.K., and B.T.M performed the research; and analyzed the data; L.C.M., J.T.K., A.J.W and R.A.M. wrote the paper.

