## Supplemental figures for "Zebrin molecular identity and cerebellar location determine Purkinje cell vulnerability in Christianson syndrome mice"

**TITLE**

**Supplemental Figures**

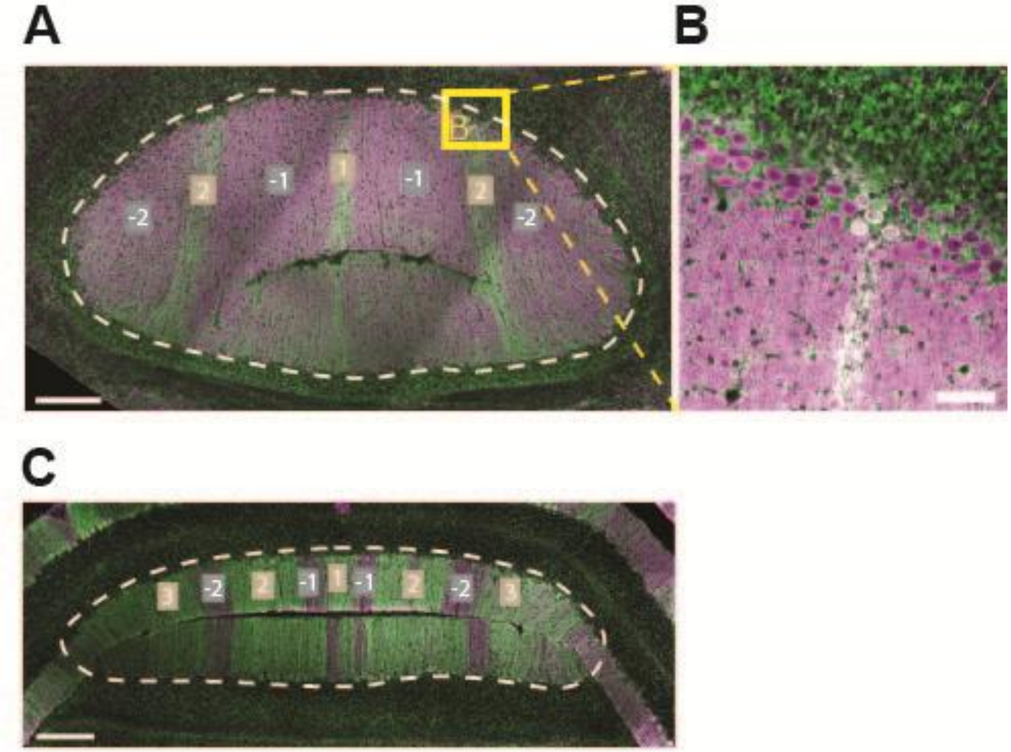


**SUPPLEMENTAL FIGURE 1. Illustrations of cerebellar cortex PC analysis.** ***(A)*** *Example image from anterior lobule III coronal section, with PCs labelled with calbindin (purple) and zebrin (green). Zebrin-negative bands -1 and -2 (grey), and zebrin-positive bands 1 and 2 (brown) are depicted. Zebrin-negative bands and PC bodies are positive for calbindin only (purple), while zebrin-positive bands and PC soma are positive for both calbindin (purple) and zebrin (green). The dashed line depicts where our analyses were taken from. PC counts were normalized to the length of the PC layer.* ***(B)*** *Higher magnification image of zoomed-in area inside yellow box in* ***(A)****, illustrating the difference between a zebrin-negative PC (purple only) and a zebrin-positive*

*PC (white due to colocalization between calbindin in purple and zebrin in green).* ***(C)*** *Example image from posterior lobule VIII/IX coronal section, with PCs labelled with calbindin (purple) and zebrin (green). Zebrin-negative bands -1 and -2 (grey), and zebrin-positive bands 1, 2, and 3 (brown) are depicted. Zebrin-negative bands and PC bodies are positive for calbindin only (purple), while zebrin-positive bands and PC soma are positive for both calbindin (purple) and zebrin (green). The dashed line depicts where our analyses were taken from. PC counts were normalized to the length of the PC layer. Scale bar, 200 µm* ***(A, C)****. Scale bar, 40 µm* ***(B).***

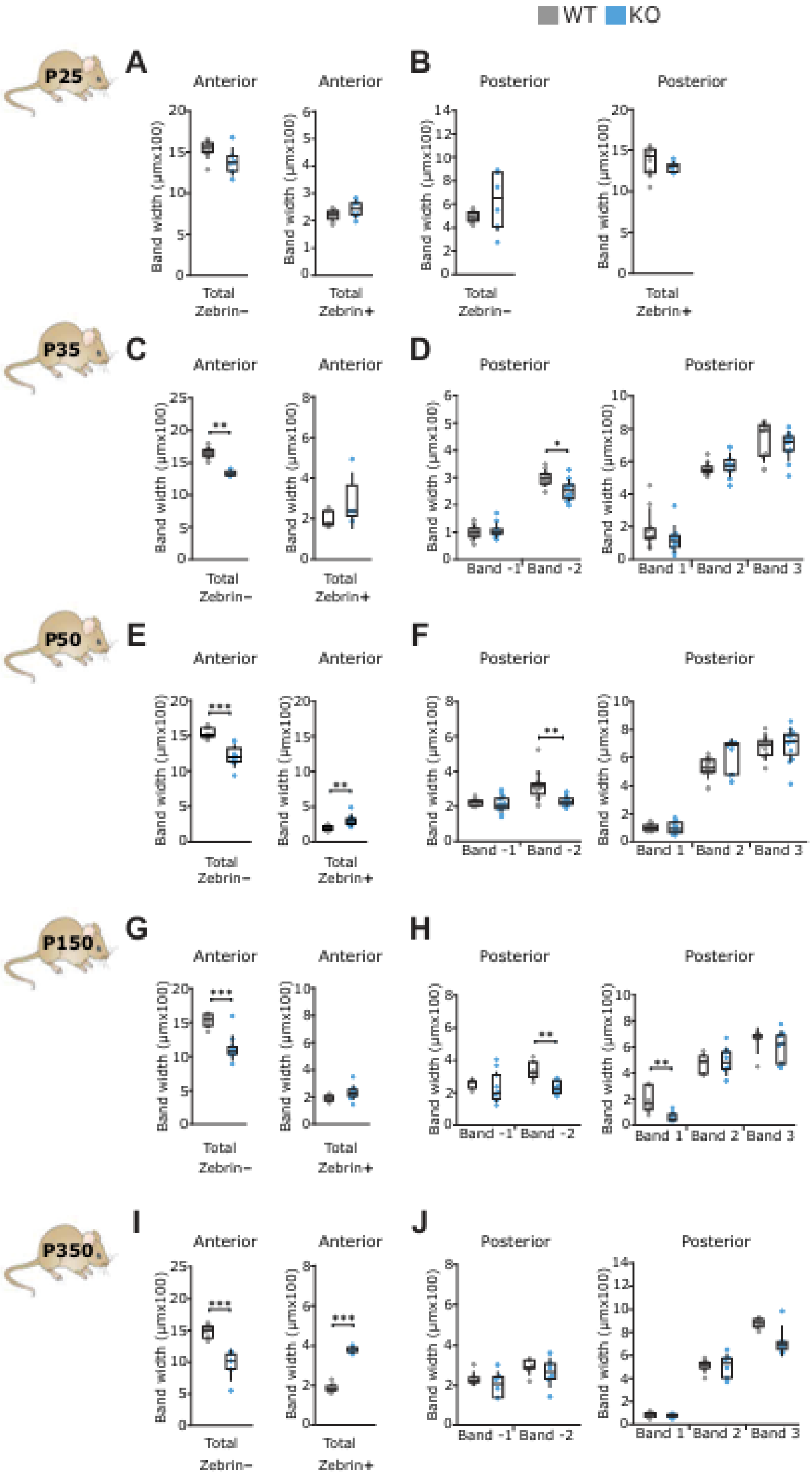
WT KO
P25
A
Anterior
Anterior
B
Posterior
Posterior
Band width (μmx100)
Total Zebrin−
Total Zebrin+
P35
C
D
**
*
Band -1
Band -2
Band 1
Band 2
Band 3
P50
E
F
***
**
P150
G
H
I
J
P350

**SUPPLEMENTAL FIGURE 2. Changes in zebrin-negative band widths are observed starting at P35.** ***(A, C, E, G, I)*** *Quantifications of the widths of the zebrin-specific bands in the anterior lobule III with the sum of the three zebrin-negative band widths shown on the left, and that of the four zebrin-positive bands shown on the right at P25* ***(A)****, P35* ***(C)****, P50* ***(E)****, P150* ***(G)*** *and P350* ***(I)****. Lower widths of zebrin-negative bands in the anterior are seen starting at p35* ***(C)****, with a possible compensatory increase in zebrin-positive band widths observed at P50* ***(E)*** *and P350* ***(I)****.* ***(B, D, F, H, J)*** *Quantifications of the widths of the zebrin-specific bands in the posterior lobule VIII with the widths of the individual 2 zebrin-negative bands shown on the left, and that of the 3 zebrin-positive bands shown on the right at P25* ***(B)****, P35* ***(D)****, P50* ***(F)****, P150* ***(H)*** *and P350* ***(J)****. A long, but transient, decrease in the zebrin-negative band width of band 2 is observed from P50-P150* ***(D, F, H)****. N=4 for WT, N=3 for CS, 2 sections per lobule per animal* ***(A-B)****; N=4 for WT, N=4 for CS, 2 sections per lobule per animal* ***(C- F)****; N=3 for WT, N=5 for CS, 2 sections per lobule per animal* ***(G-H)****; N=4 for WT, N=3 for CS, 2 sections per lobule per animal* ***(I-J)****. For statistics, student's t-test for normally distributed data and Mann-Whitney U test for non-normally distributed data were used; *P<0.05, **P<0.01, ***P<0.001; n.s. refers to P>0.05.*

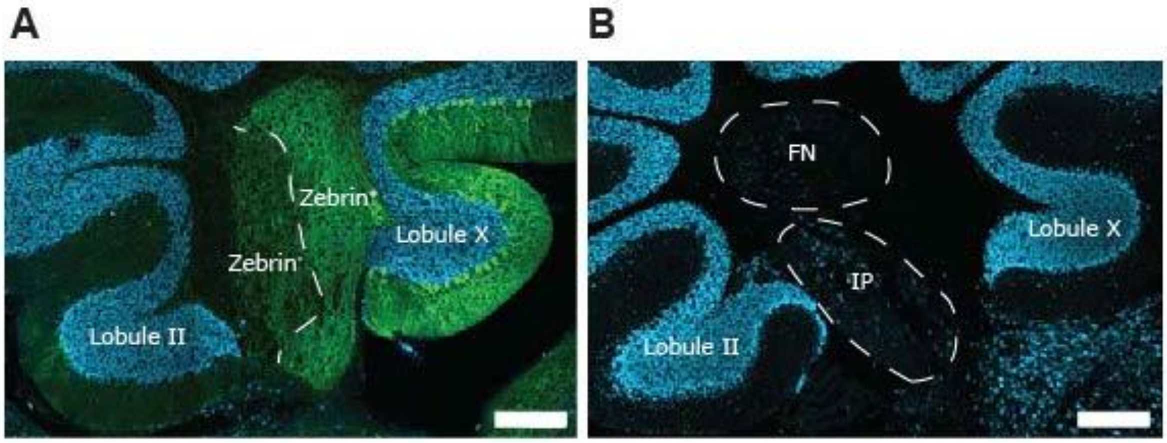


**SUPPLEMENTAL FIGURE 3. Illustration of CN analyses.** ***(A)*** *Representative image of the CN, with the dashed line showing zebrin-negative and zebrin-positive PC input delineation in the CN. Zebrin-negative area is adjacent to lobule II and the anterior cerebellar cortex, while zebrin-positive area is adjacent to lobule X and the posterior cerebellar cortex. Analyses in the CN were based on this delineation, with the anterior CN region referring to the zebrin-negative side, and the posterior CN region referring to the zebrin-positive side.* ***(B)*** *Representative image of the CN, with the dashed circles showing the fastigial (FN) and interposed (IP) nuclei regions. Analyses of CN morphology, both area and CN soma count, were taken in these respective regions. Scale bar, 200 µm* ***(A, B).***

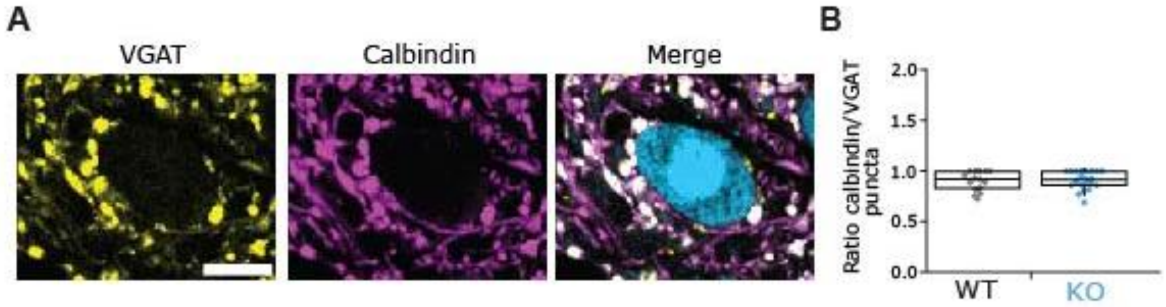


**SUPPLEMENTAL FIGURE 4. Validation of calbindin as a marker for functional presynaptic GABAergic PC terminals onto large cells of the CN.** ***(A)*** *Representative images of PC input onto a large CN cell, labeled with VGAT (yellow) and calbindin (purple).* ***(B)*** *VGAT-positive puncta and calbindin-positive puncta show very high colocalization, both in WT (grey) and CS (blue) cells, as evidenced by no significant differences between both distributions, thus supporting the use of calbindin-positive input onto CN cells as a representative marker of functional presynaptic GABAergic terminals. N=3 animals for WT, N=3 animals for CS, two sections per animal. For statistics, Student's t-test for normally distributed data and Mann Whitney U test for non-normally distributed data were used; n.s. refers to P>0.05. Scale bar, 10 µm.*

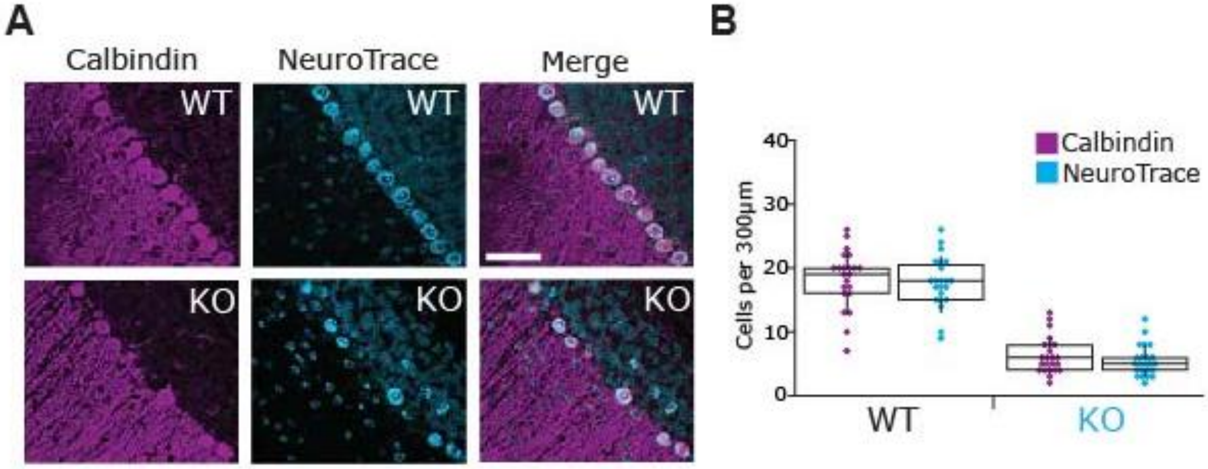


**SUPPLEMENTAL FIGURE 5. Validation of calbindin as a marker for PCs using a fluorescent Nissl stain.** ***(A)*** *Representative images taken from coronal anterior lobule III in age P50 animals showing immunohistochemistry for calbindin (purple) and fluorescent Nissl stain NeuroTrace (blue), for WT (top) and CS (bottom) mice.* ***(B)*** *Quantification of the density of calbindin (purple) and NeuroTrace (blue) cells per 300 µm sections, showing WT (left) and CS (right) distributions. Manual identification of calbindin-positive and/or NeuroTrace-positive PCs revealed no significant differences in labeling, neither in WT nor CS. Statistically, all manually identified calbindin-positive PCs were also NeuroTrace-positive. With apparent PC loss in CS at this age, using calbindin as a marker for PCs appears to be equally reliable as using NeuroTrace staining for neuronal nuclei. N=3 animals for WT, N=3 animals for CS, two sections per animal. For statistics, Student's t-test for normally distributed data and Mann Whitney U test for non-normally distributed data were used; n.s. refers to P>0.05. Scale bar, 60 µm.*

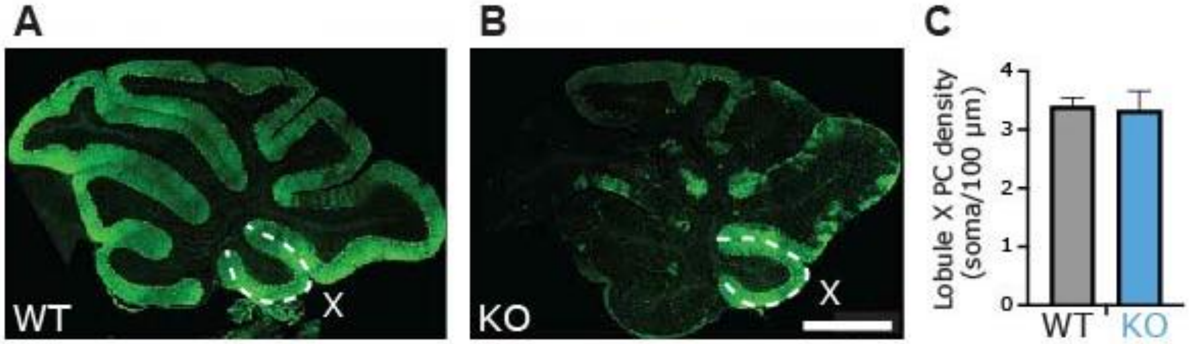


**SUPPLEMENTAL FIGURE 6. PC density in lobule X is unchanged in CS compared to WT at P350.** ***(A-B)*** *Representative image of a sagittal cerebellar slice from a WT* ***(A)*** *and a CS* ***(B)*** *mouse showing immunohistochemistry for calbindin (green) to stain PCs.* ***(C)*** *Quantification of PC density in lobule X per 100µm showing average density for WT (grey) and CS (blue). No differences were found between WT and CS mice PC density. N=4 for WT and N=4 for CS. For statistics, Student's t-test was performed n.s. refers to P>0.05. Scale bar, 400µm.*
